# A striatal interneuron circuit for continuous target pursuit

**DOI:** 10.1101/386490

**Authors:** Namsoo Kim, Haofang E. Li, Ryan N. Hughes, Glenn D. R. Watson, David Gallegos, Anne E. West, Il Hwan Kim, Henry H. Yin

**Author notes:** Correspondence should be addressed to Henry Yin, Box 91050, Duke University, Durham, NC 27708.

## Abstract

Most adaptive behaviors require precise tracking of targets in space. In pursuit behavior with a moving target, mice use distance to target to guide their own movement continuously. Here we show that in the sensorimotor striatum, parvalbumin-positive fast-spiking interneurons (FSIs) can represent the distance between self and target during pursuit behavior, while striatal projection neurons (SPNs), which receive FSI projections, can represent self-velocity. FSIs are shown to regulate velocity-related SPN activity during pursuit, so that movement velocity is continuously modulated by distance to target. Moreover, bidirectional manipulation of FSI activity can selectively disrupt performance by increasing or decreasing the self-target distance. Our results reveal a key role of the FSI-SPN interneuron circuit in pursuit behavior, and elucidate how this circuit implements distance to velocity transformation required for the critical underlying computation.

## Introduction

Whether pursuing a prey or approaching a mate, natural behaviors often involve continuous tracking of targets in space. Yet the neural substrates of such pursuit behavior remain poorly understood. Technical limitations have prevented the study of natural pursuit behavior in freely moving animals^1–4^. In this study we designed a new behavioral task for freely moving mice. In this task, mice follow a continuously moving target that delivers sucrose reward. Using 3D motion capture, we were able to track not only the position of the animal but also the distance to target, a crucial variable for accurate pursuit. This task allows us to compare, for the first time, continuous behavioral variables and neural activity recorded at the same time in freely moving animals.

We studied the contribution of the inhibitory interneuron circuit in the sensorimotor striatum to pursuit behavior. The striatum is a major basal ganglia (BG) nucleus that has been implicated in motor control, compulsive behavior, and habit formation ^5–9^. A critical circuit in the striatum is formed by the parvalbumin positive GABAergic fast-spiking interneurons (FSIs) and striatal projection neurons (SPNs)^10, 11^. FSIs, which constitute less than 1% of the striatal neuronal population, receive glutamatergic inputs from the cerebral cortex and project to many SPNs, which make up over 90% of the population ^12, 13^. The sensorimotor striatum is characterized by the highest expression of FSIs. Reduced numbers of FSIs is associated with neuropsychiatric disorders such as Tourette’s syndrome and obsessive-compulsive disorder ^14, 15^. Behavioral studies have also implicated the FSIs in choice behavior and habitual lever pressing ^13, 16^.

Recent work has shown that sensorimotor SPN activity is often highly correlated with movement velocity ^17, 18^, and stimulation of striatal output alters movement velocity frequency-dependently ^19^. Yet it remains unclear how, during target-driven pursuit behavior, the striatal output is commanded by representation of distance to target. We hypothesize that the FSI-SPN interneuron circuit plays a critical role in pursuit behavior, allowing cortical inputs representing the spatial relationship between self and target to reach the BG. In particular, the distance representation can be converted into instantaneous velocity commands during pursuit behavior. To test this hypothesis, we used wireless *in vivo* electrophysiology, calcium imaging, and cell-type specific manipulation of neural activity to examine the contributions of the striatal microcircuit to pursuit behavior in freely moving mice. Our results established for the first time a key role of this local striatal circuit in pursuit behavior.

## Results

### Behavioral task

We recorded head and target positions during pursuit behavior using motion capture at 100 frames per second (**Fig. 1a-b**). All mice rapidly learned to follow the moving target (**Extended Data Fig. 1, Video 1**) and showed comparable performance with constant target velocity (16 mm/s, leftward or rightward) and variable velocity (5-48 mm/s). Because in our task pursuit behavior is self-initiated, we separated periods in which the mice followed the target and periods in which they did not. ‘Following’ is defined as staying in a spatially defined zone close to the target for at least 800 ms (<15 mm x-axis, <10 mm in y axis, <20 mm in z axis), when the head is next to the sucrose spout. Otherwise, behavior is classified as ‘not following’ (**Fig. 1b-d**; for variable velocity see **Extended Data Fig. 1**). When the animal is not following, it could be engaged in any number of activities, such as grooming, orienting, rearing, and so on. A cross-correlation analysis suggests that mice use distance information to adjust their velocity (**Fig. 1e**). With distance as the reference variable, a positive lag means that distance leads velocity, and that the distance representation is used to guide velocity. This is exactly what we found. This lag measure is also positively correlated with the distance, the key measure of pursuit performance: the worse the pursuit performance, the more self-velocity lags self-target distance, as expected if distance is used to adjust velocity.

**Fig. 1.**
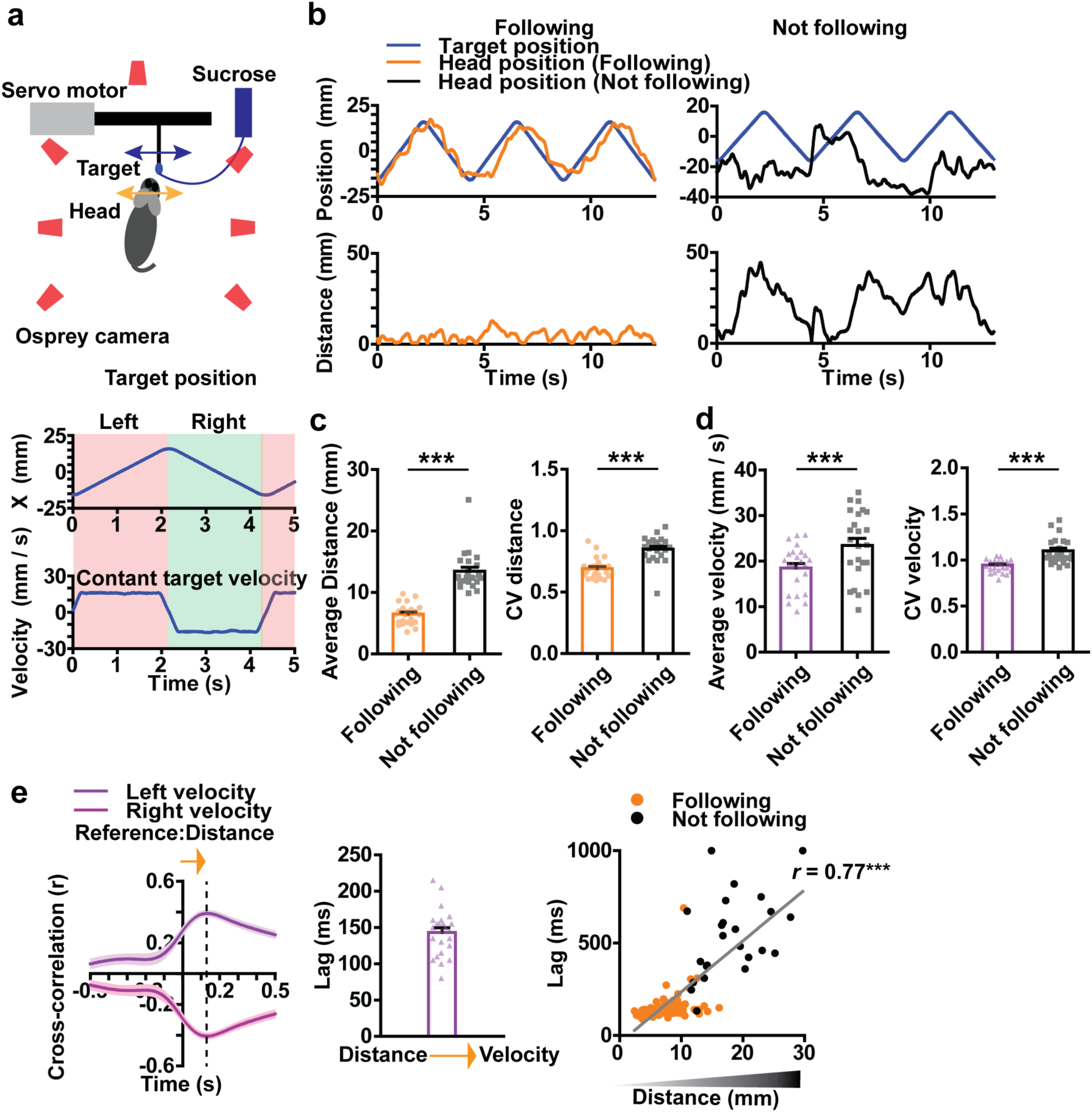
Summary of behavioral results. a) Illustration of the pursuit task and location of the infrared cameras for 3D motion capture. The target moves continuously from side to side, each cycle consisting of both leftward and rightward movements. During pursuit, the target could be on either side of the mouse. Right, position and velocity traces of the target, which moves along the x-axis at a constant velocity of 16 mm/s (see **Extended Fig. 1** for variable velocity condition). The total distance traveled from left to right is 32 mm. Thus, it takes the target ∼2 seconds to move from one extreme to another. b) Representative traces of mouse head and target position. Left panel shows the head and target positions during ‘Following’. ‘Following’ is defined as a period in which the head is close to the target (<15 mm x-axis, <10 mm in y axis, <20 mm in z axis). Right panel shows the head and target positions during ‘Not following’. Also see **Videos 1 and 2** for examples. c) Average distance is significantly reduced during following (paired t-test, p < 0.0001). Error bars indicate ± s.e.m. *** p < 0.0001. Coefficient of variation (CV) in distance shows that variance is also increased when animals are not following. d) Average self-velocity is significantly lower during following (paired t-test, p < 0.0001). CV in velocity is also increased when animals are not following. e) Illustration of the temporal relationship between distance and self-velocity. Cross correlation analysis reveals the lag between these two variables, showing that distance leads self-velocity during pursuit behavior. In addition, the lag shows a positive correlation with distance (p < 0.0001). When the mouse is actively pursuing and maintaining a small distance, the lag between distance and velocity is small, but the lag increases as pursuit performance declines.

### Neuronal representations of distance and velocity

To elucidate how striatal neurons contribute to pursuit behavior, we recorded striatal neurons while mice performed the task (453 neurons from 24 mice). We classified FSIs and SPNs based on their different average firing rates and spike waveforms. We identified 362 putative SPNs in the dorsolateral striatum (trough to peak width: 570.8 ± 2.3 µs, firing rate: 2.1 ± 0.1 Hz) and 91 FSIs (width: 173.9 ± 4.9 µs, firing rate: 20.4 ± 0.9 Hz, **Extended Data Fig. 2**). The proportion of FSIs is higher than expected from anatomical studies (1-2%) because these neurons have much higher firing rates compared to SPNs. Our cell type classification is confirmed by optogenetic tagging of PV+ FSIs.

FSIs and SPNs showed distinct representations of behavioral variables: FSIs more commonly represented self-target distance (56%) whereas SPNs (32%) more commonly represented velocity (chi square = 88.71, *P* <0.0001, **Tables 1** and **2**, **Extended Fig. 3**). As shown in **Fig. 2**, there are two distinct populations of SPNs: One population increases firing as leftward velocity increases (**Fig. 2a-c)** and the other increases firing as rightward velocity increases (**Fig. 2d-f).** When we analyzed the relationship between single unit activity and velocity from the entire session, the correlation coefficient is high. By comparison, correlation with other variables is much lower (**Fig. 2g, Extended Data Fig. 4**). Using the population average activity as a measure of the ensemble activity, there is strong correlation between neural activity and velocity **(Fig. 2h)**. Activity quickly decreases and reaches a floor during movement in the opposite direction. Overall, the relationship between neural activity and velocity can be described with a sigmoidal function ^20^.

**Table 1.** SPN electrophysiology summary.

|  | Distance (n) | Velocity (n) | Acceleration (n) | Reward (n) | Unclassified (n) | Total (n) |
| --- | --- | --- | --- | --- | --- | --- |
| # neurons | 5.2% (19) | 32.0% (116) | 3.3% (12) | 0.0% (0) | 59.4% (215) | 100.0% (362) |

|  | Left striatum (n) | Right striatum (n) |
| --- | --- | --- |
| Left distance | 2.3 % (5) | 1.4 % (2) |
| Right distance | 3.2 % (7) | 3.6 % (5) |
| Left velocity | 19.8 % (44) | 10.7 % (15) |
| Right velocity | 11.3 % (25) | 22.9 % (32) |
| Left acceleration | 1.4 % (3) | 1.4 % (2) |
| Right acceleration | 2.3 % (5) | 1.4 % (2) |
| Reward | 0.0% (0) | 0.0% (0) |
| Unclassified | 59.9 % (133) | 58.6 % (82) |
| Total | 100.0 (222) | 100.0 (140) |

**Table 2.** FSI electrophysiology summary.

|  | Distance (n) | Velocity (n) | Acceleration (n) | Reward (n) | Unclassified (n) | Total (n) |
| --- | --- | --- | --- | --- | --- | --- |
| # neurons | 56 % (51) | 11.0 % (10) | 5.5 % (5) | 0.0% (0) | 27.5 % (25) | 100.0% (91) |

|  | Left striatum (n) | Right striatum (n) |
| --- | --- | --- |
| Left distance | 22.9 % (11) | 20.9 % (9) |
| Right distance | 20.8 % (10) | 48.8 % (21) |
| Left velocity | 10.4 % (5) | 4.7 % (2) |
| Right velocity | 2.1 % (1) | 4.7 % (2) |
| Left acceleration | 2.1 % (1) | 0.0 % (0) |
| Right acceleration | 6.3 % (3) | 2.3 % (1) |
| Reward | 0.0 % (0) | 0.0 (0) |
| Unclassified | 35.4 % (17) | 18.6 % (8) |
| Total | 100.0 (48) | 100.0 (43) |

**Fig. 2.**
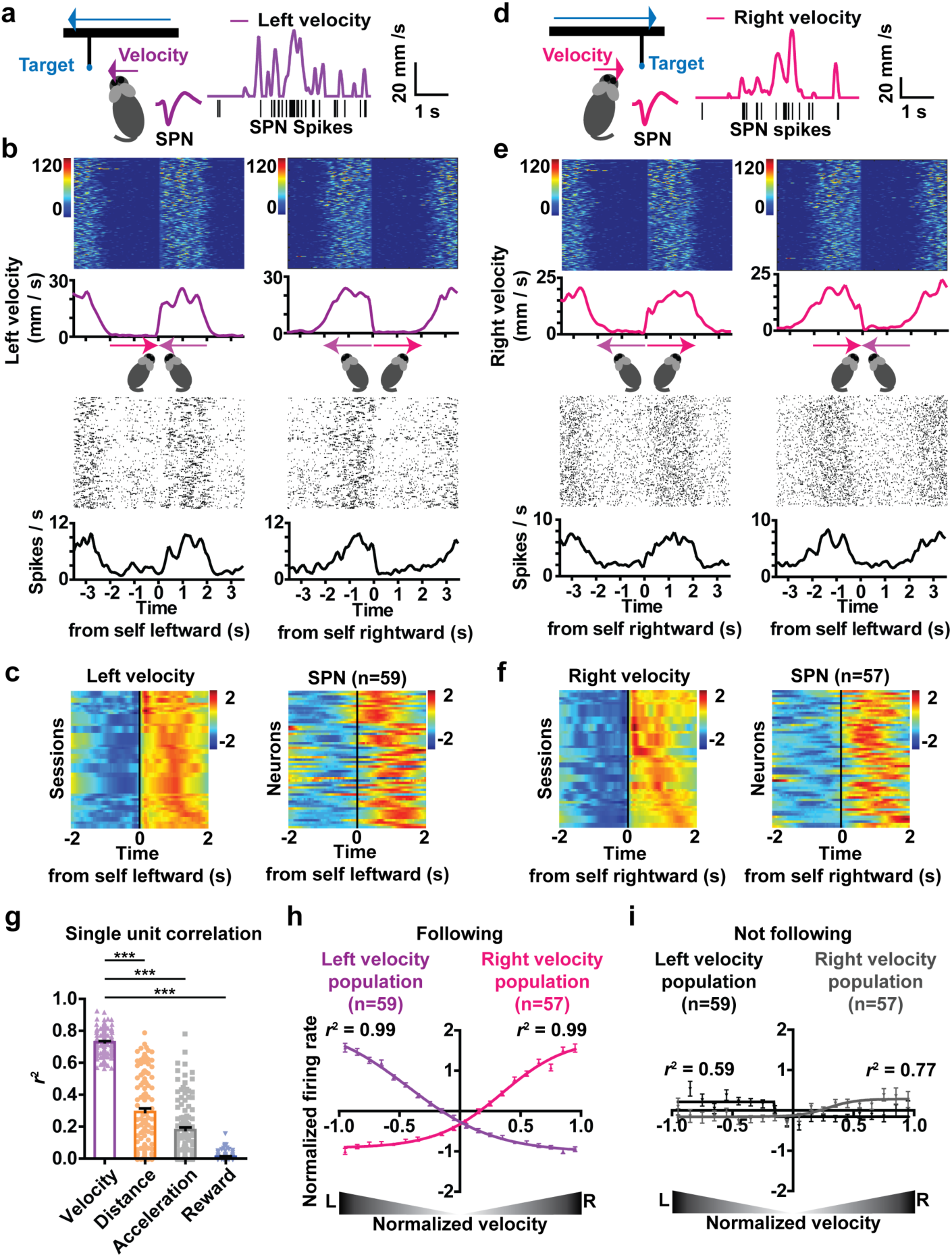
SPNs represent self velocity when animal is following target. a) A representative SPN and spike waveform. The firing rate of this neuron is positively correlated with leftward velocity. b) Raster plots comparing single unit activity of a representative left velocity SPN and velocity. Time zero is the start of leftward or rightward pursuit behavior. Note the similarity between velocity and firing rate. c) Spike density function of all left velocity SPNs (z-scored values). Each row represents average velocity measure and the firing rate of corresponding SPNs from a single session. Note that some rows in the velocity plot are identical, because multiple neurons from the same mouse are correlated with the same velocity variable. d) A representative SPN and spike waveform. The firing rate of this neuron is positively correlated with rightward velocity. e) Raster plots comparing velocity and firing rates of a right velocity SPN. f) Spike density function of all right velocity SPNs (z-scored values). g) The correlation coefficient between firing rate of individual velocity SPNs and different behavioral variables. In each case, velocity has the highest r^2^ value (repeated measures ANOVA: main effect of variable, F _3,463_ = 453.8, p < 0.0001; Bonferroni post hoc: velocity vs. distance p < 0.0001, velocity vs. acceleration p < 0.0001, velocity vs. reward p < 0.0001). Error bars indicate ± s.e.m. *** p < 0.0001. h) Population average activity of all velocity-correlated SPNs. i) Velocity SPNs show much weaker correlation if the mouse is not following.

Our results suggest that the ensemble activity of many neurons with similar velocity representations (e.g. leftward) is used as the signal for velocity command. With negative feedback control, the actual velocity achieved closely matches the command. Interestingly, the correlation with velocity is significantly reduced when the mice are not following, for both populations of velocity-correlated neurons (**Fig. 2i**). This finding suggests that the relationship between velocity and SPN output is contingent upon self-initiated pursuit behavior

On the other hand, many FSIs showed continuous representation of target distance (**Fig. 3, Extended Data Fig. 5**). Their firing rates varied monotonically with target distance. The representation of target distance is independent of target or self-movement direction (whether leftward or rightward). One group increases firing as left distance increases (**Fig. 3a-c**), whereas a second group increases firing as right distance increases (**Fig. 3d-f**). By comparison, the correlation with other variables is much lower (**Fig. 3g**). There is a strong linear relationship between the population activity of distance FSIs and distance (**Fig. 3h**). For both populations, the mean firing rate corresponds to roughly the ‘straight ahead’ condition. For the ‘left+’ neurons, the highest firing rate is reached at the leftmost extreme and lowest firing rate at the rightmost extreme, and vice versa for the ‘right+’ neurons. **Fig. 3i** shows that the correlation with distance is significantly reduced when the mice are not following.

**Fig. 3.**
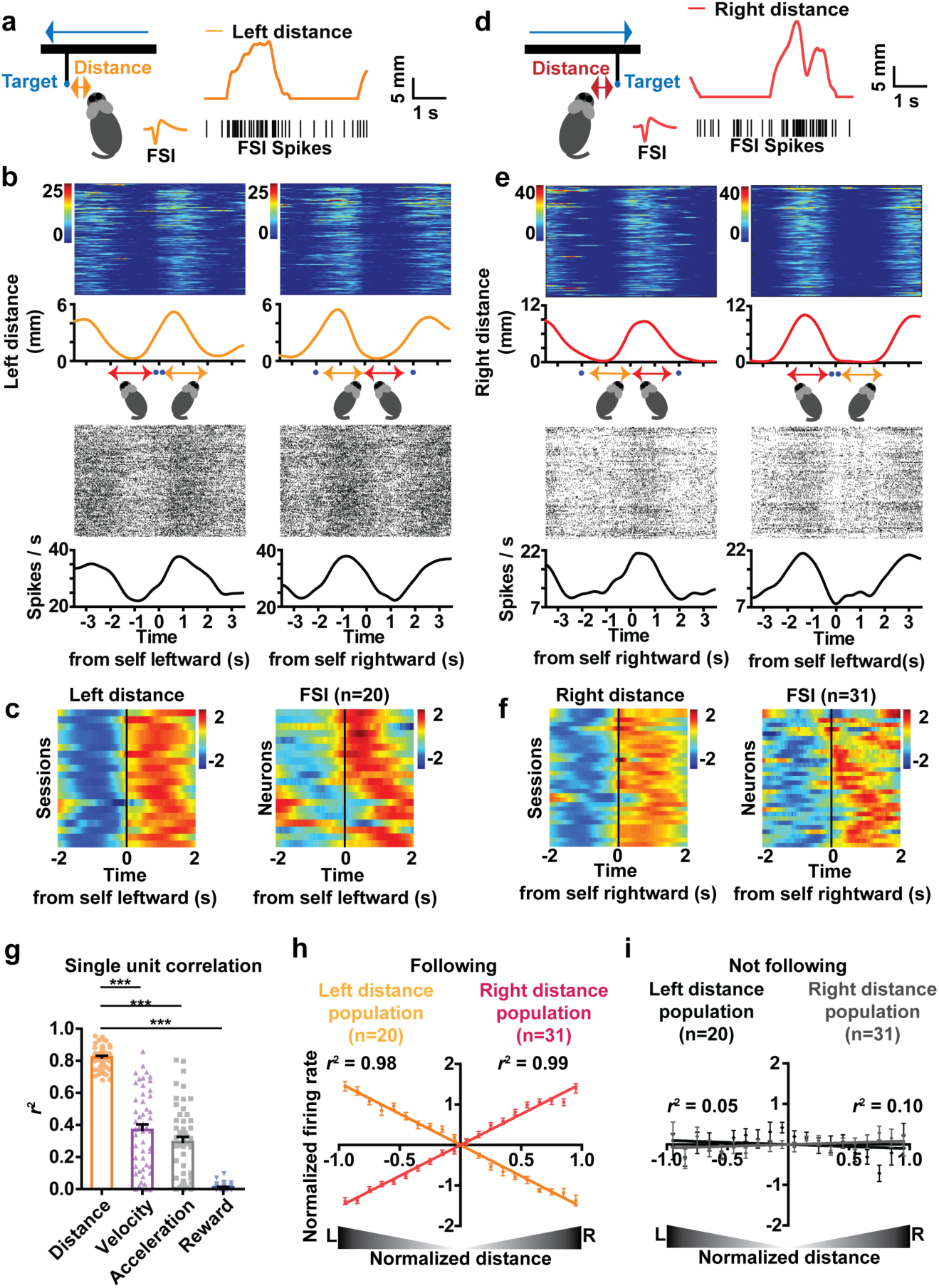
FSIs can represent distance to target. a) A representative FSI and spike waveform. The firing rate of this neuron is positively correlated with distance to target on the left side of the head, regardless of whether the mouse is moving leftward or rightward. b) Raster plots comparing single unit activity of a representative left distance FSI and distance. Time zero is the start of leftward or rightward pursuit. c) Spike density function of all left distance FSIs. Each row represents average distance measure and the firing rate measure of corresponding FSI from a single session. Note that some rows in the velocity plot are identical, because multiple neurons are correlated with the same distance variable. d) A representative FSI and spike waveform. The firing rate of this neuron is proportional to distance to target on the right side of the head, regardless of whether the mouse is moving leftward or rightward. e) Raster plots comparing single unit activity of a representative right distance FSI and distance. Time zero is the start of leftward or rightward pursuit. f) Spike density function of all right distance FSIs. g) The correlation coefficients between firing rate of distance FSIs and alternative behavioral variables. In each case, distance has the highest coefficient of correlation with firing rate (repeated One-way ANOVA F _3,203_ = 193.0, p < 0.0001; Bonferroni post-hoc: distance vs. velocity p < 0.0001, distance vs. acceleration p < 0.0001, distance vs. reward p < 0.0001). Error bars indicate ± s.e.m. *** p < 0.0001. h) Population average activity of all distance-correlated FSIs. i) Distance FSIs show much weaker correlation when not following.

Because the striatum contains many different types of neurons, classification of a relatively rare class like the FSI based purely on firing rate and spike waveform may not be reliable. There are other interneurons that also have higher firing rates than SPNs ^12^. To confirm our classification, we used optrodes to selectively stimulate and record PV+ FSIs at the same time. We injected a Cre-dependent channelrhodopsin (DIO-ChR2) in transgenic mice that express Cre-recombinase in PV+ neurons (PV-Cre)^21^. Only neurons that are activated by light stimulation with a short latency (< 8 ms) are classified as FSIs (**Table 3**). The optogenetically tagged cells show similar waveforms and firing rates as those we classified as FSIs (**Fig. 4a-b**). More importantly, most identified FSIs are also correlated with distance to target (**Fig. 4c-e**). The population average activity shows a highly linear relationship with distance (**Fig. 4f**).

**Table 3.** FSI optotagging summary.

|  | Distance (n) | Velocity (n) | Acceleration (n) | Reward (n) | Unclassified (n) | Total (n) |
| --- | --- | --- | --- | --- | --- | --- |
| # neurons | 73.3 % (11) | 6.7 % (1) | 0.0 % (0) | 0.0 % (0) | 20.0 % (3) | 100.0% (15) |

**Fig. 4.**
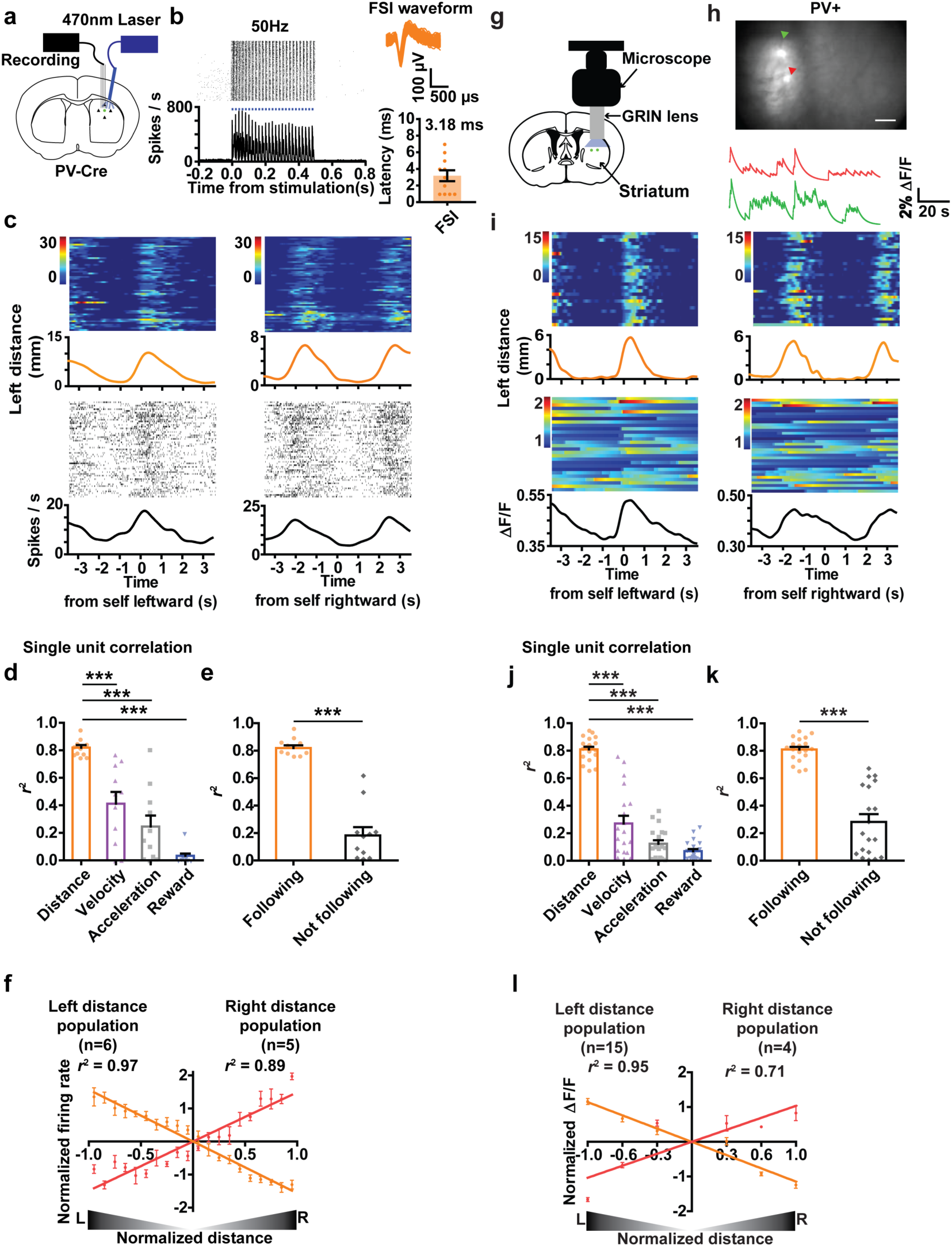
Optotagging single unit activity and in vivo calcium imaging of FSI activity during pursuit. a) Schematic illustration of simultaneous optogenetic stimulation and recording using a chronically implanted optrode. An AAV vector with Cre-inducible ChR2 is injected into the sensorimotor striatum of PV-Cre mice to allow selective activation of PV+ FSIs. b) A representative left distance neuron that is optically tagged. ChR2-induced stimulation at 50 Hz reliably triggered spiking in FSIs with a short latency (average latency = 3.18 ± 0.65 ms). c) Representative left distance FSI that is optically tagged. The firing rate of this neuron is proportional to leftward distance. Time zero is the start of leftward or rightward pursuit. d) Correlation between optotagged distance FSIs and different behavioral variables (repeated One-way ANOVA F _5, 43_ = 30.98, p < 0.0001; Bonferroni post-hoc: distance vs. velocity p < 0.0001, distance vs. acceleration p < 0.0001, distance vs. reward p < 0.0001). Error bars indicate ± s.e.m. *** p < 0.0001. e) Correlation between FSI distance neurons and distance is significantly lower when the animal is not following the target (p < 0.0001). f) Population summary of all optotagged distance FSI neurons. g) Calcium imaging with an implantable GRIN lens (1.8 mm diameter). Schematic illustration of the GRIN lens implant. A Cre-dependent GCamp6s virus was injected into the sensorimotor striatum of PV-Cre mice. h) Representative image of FSIs. Red and green traces indicate two simultaneously imaged FSIs. Because FSIs are rare, only 2-11 cells were recorded per mouse. Also see **Video 3**. i) Representative left distance FSI. Calcium signal from this neuron is proportional to leftward distance. Time zero is the start of leftward or rightward pursuit. j) Correlation between distance FSIs and different behavioral variables (repeated One-way ANOVA, F *_3, 75_* = 103.4, p < 0.0001; Bonferroni post hoc analysis: distance vs. velocity p < 0.0001; distance vs. acceleration p < 0.0001; distance vs. reward p < 0.0001). k) Correlation between FSI distance neurons and distance is significantly lower when the animal is not actively following (paired t-test, p < 0.0001). l) Population summary of all distance FSIs.

Finally, to confirm our *in vivo* electrophysiological results, we also utilized calcium imaging with miniaturized endoscopes to record FSI activity (**Fig. 4g-h, Table 4**)^22, 23^. In agreement with our electrophysiological data, we found many FSIs whose activity represents distance (**Fig. 4i-k**). Population average activity of FSIs also showed a linear relationship with distance to target, with two opponent populations (**Fig. 4l**).

**Table 4.**
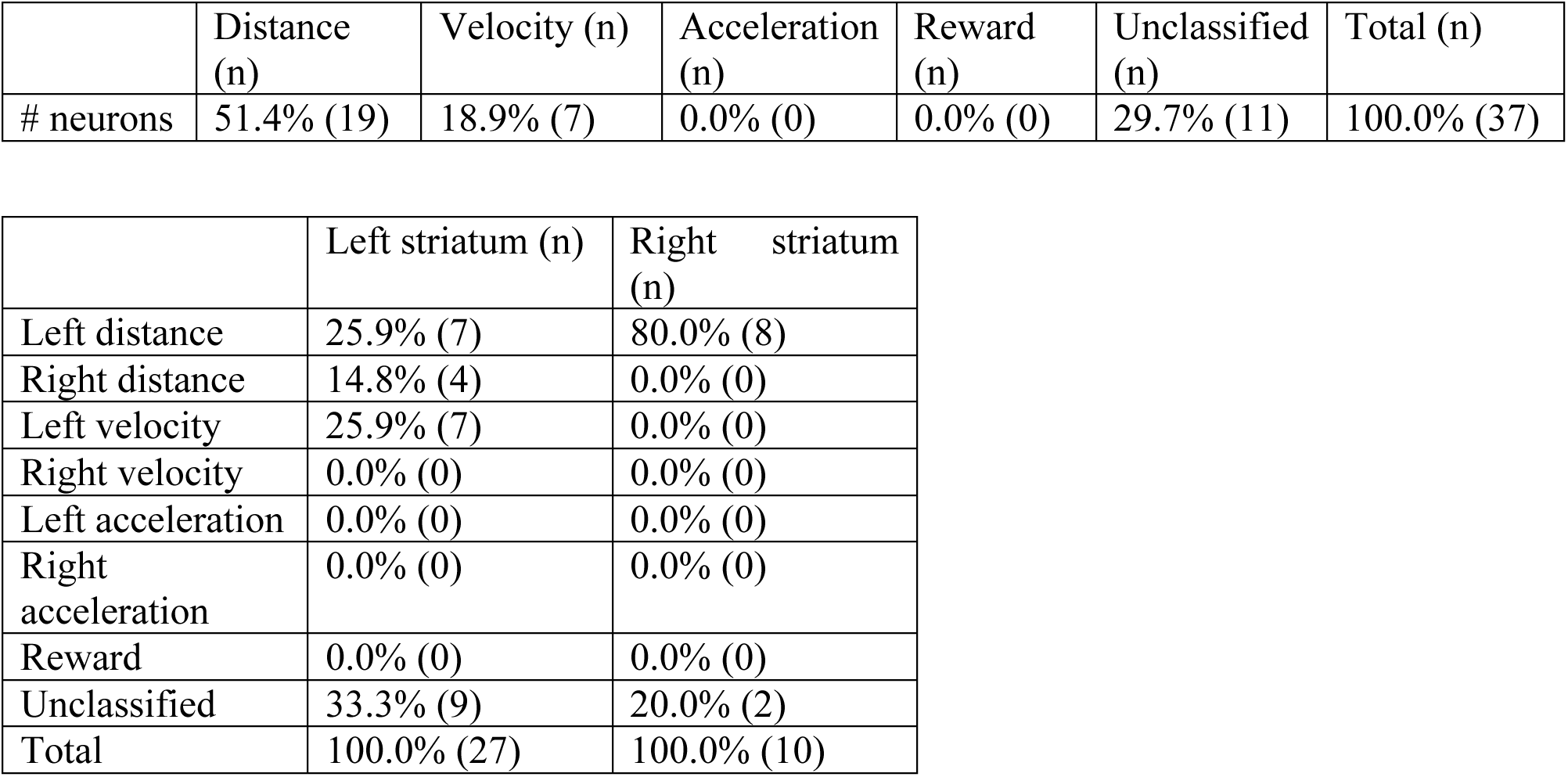
Calcium imaging summary.

### Manipulations of neural activity

Our electrophysiological and imaging results show that distance to target, the key variable in pursuit behavior, is represented by striatal FSI neurons. However, they do not tell us whether this representation is required for successful pursuit. To test necessity, we silenced FSIs using the tetanus toxin light chain (TeLC). TeLC prevents neurotransmitter release by cleaving its vesicle associated membrane protein 2 ^24, 25^. We used Cre-dependent TeLC (AAV-flex-TeLC) in PV-Cre mice (**Fig. 5a**), thus expressing TeLC in a cell type- and region-specific manner (**Fig. 5b, Extended Data Fig. 6**). TeLC was injected after training, and we tested the mice again following recovery. FSI silencing markedly disrupted target pursuit performance, without affecting overall movement velocity during the session (**Fig. 5c-e**).

**Fig. 5.**
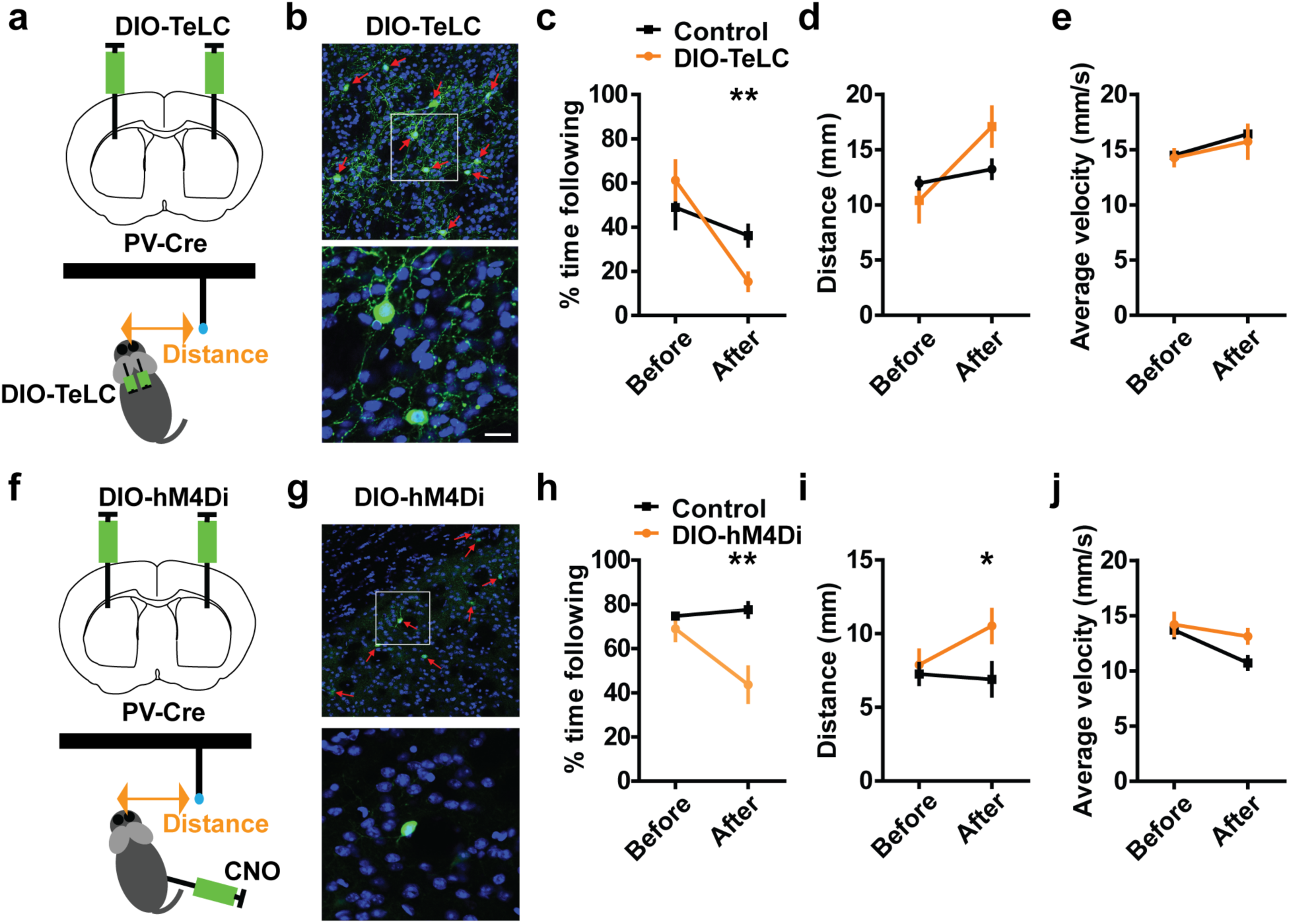
Silencing FSI activity disrupts pursuit performance. a) To silence neural activity in FSIs, we injected the Cre-dependent DIO-TeLC virus into the striatum of PV-Cre mice. TeLC silences neurons by cleaving the vesicular associated membrane protein 2 and preventing transmitter release b) GFP staining showing FSIs infected with GFP-tagged TeLC, (control: n = 6, DIO-TeLC: n = 7). c) Duration of self-initiated pursuit behavior (% time spent following) is significantly reduced by TeLC. A repeated measures ANOVA revealed a significant interaction: F _1, 11_ = 6.88, p = 0.024. Post hoc test showed a significant effect of TeLC (p = 0.0051). Error bar indicates ± s.e.m. ** P < 0.01. d) Distance error is greater in the TeLC group compared to the Control group (GFP only). Repeated measure ANOVA: Interaction: F _1, 11_ = 5.92, P < 0.05. Post hoc unpaired t-test, p = 0.09. e) TeLC did not affect velocity: no significant interaction: F _1, 11_ = 0.08, P = 0.77, no effect of Time, F _1, 11_= 12.73, P = 0.05, or of Group: F _1, 11_= 0.94, P = 0.67. f) To reversibly inactivate FSIs, we injected the Cre-dependent DIO-hM4Di virus into the striatum. g) GFP staining showing FSIs in the striatum infected with GFP-tagged DREAAD (control: n = 6, DIO-hM4Di: n = 6). h) hM4Di reduced % time spent following the target. There was a significant interaction: F _1, 10_ = 15.2, P = 0.0030. Postdoc test revealed a significant effect of CNO after injection (P = 0.0028). **P < 0.01. Error bar indicates ± s.e.m. i) hM4Di increased distance: Interaction: F _1, 10_ = 9.26, P = 0.044. * P < 0.05. j) hM4Di did not affect velocity. No interaction: F _1, 10_ = 5.11, P = 0.12, main effect of time: F _1, 10_ = 23.21, P < 0.01, no effect of Group: F _1, 10_= 12.09, P = 0.11).

TeLC effects are permanent and it can be difficult to rule out compensatory plasticity due to reduced FSI synaptic transmission. We therefore used designer receptors exclusively activated by designer drugs (DREADDs) to suppress FSI activity reversibly (**Fig. 5f**) ^26^. We injected an AAV vector encoding a Cre-dependent designer G protein-coupled hM4Di receptor bilaterally (**Fig. 5g, Extended Data Fig. 7**). Injection of the synthetic ligand clozapine-N-oxide (CNO) for the hM4Di receptor reduced the amount of time spent following the target (**Fig. 5h**) and increase distance to target (**Fig. 5i**) without affecting movement velocity in general (**Fig. 5j**).

To test the effects of temporally specific manipulation of FSI activity, we performed optogenetic experiments to manipulate FSI activity bidirectionally during behavior ^21, 27^. FSIs were excited using the excitatory channelrhodopsin (ChR2) injected in the sensorimotor striatum of PV-Cre mice (**Fig. 6a, Extended Data Fig. 8**) and inhibited using the inhibitory channelrhodopsin stGtACR2, which conducts anions (both can be activated by blue light, **Fig. 6b, Extended Data Fig. 9**) ^28–30^. As expected, bilateral ChR2 activation of FSIs increased distance to target, whereas bilateral stGtACR2 inhibition of FSIs decreased distance (**Fig. 6c, Extended Data Fig. 10**). Since FSIs are known to provide inhibition of SPNs, these manipulations have the opposite effects on velocity (**Fig. 6d**).

**Fig. 6.**
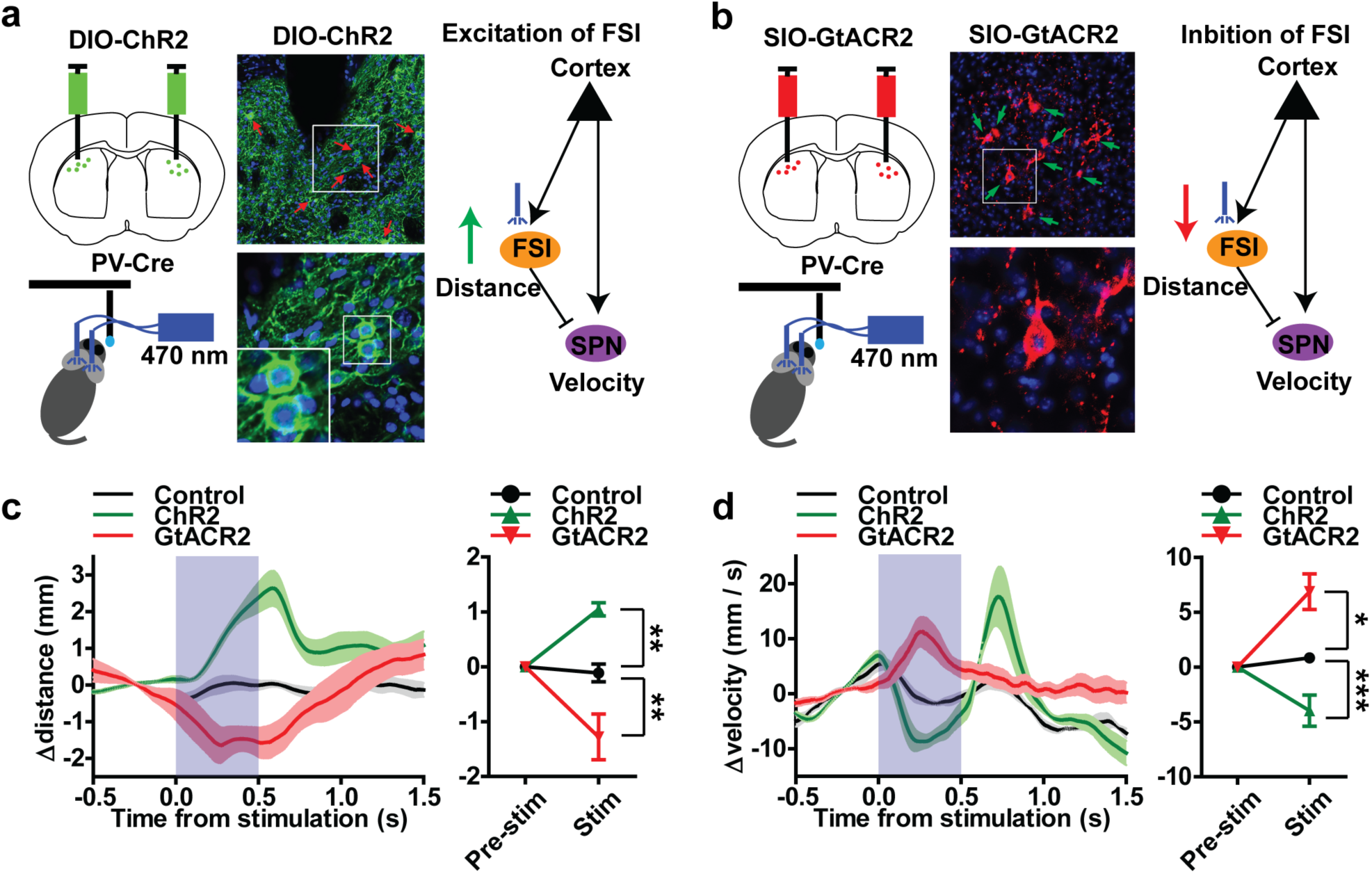
Optogenetic manipulation of FSI activity disrupts pursuit performance. a) We injected Cre-dependent excitatory channelrhodopsin (AAV-EF1a-DIO-ChR2-EYFP) into the sensorimotor striatum of PV-Cre mice. Middle, GFP staining showing FSIs infected with ChR2. This manipulation is expected to increase FSI activity and reduce target SPN activity. b) We injected Cre-dependent inhibitory channelrhodopsin (AAV-hsyn1-SIO-GtACR2-Fusion Red) to reduce FSI activity and increase target SPN activity via disinhibition. c) Effect of photo-stimulation on self-target distance. Increasing FSI activity with ChR2 increased distance, whereas decreasing FSI activity with GtACR2 decreased distance. (repeated measures two-way ANOVA, Interaction: F _2, 11_ = 12.68, p < 0.0014, Group: F _2, 11_ = 12.68, p = 0.0014; Time: F _1, 11_ = 0.35, p = 0.57, Boferroni post-hoc: control vs. ChR2, p < 0.01; control vs. GtACR2, p < 0.01). PV-Cre control: n = 4; ChR2: n = 4; GtACR2: n = 6. Error bar indicates ± s.e.m. ** P < 0.01. d) Effect of photo-stimulation on self velocity. Since FSIs inhibit SPNs, the effects on velocity are expected to be the opposite as those on distance in c. ChR2 decreased velocity, whereas GtACR2 increased velocity (repeated measures two-way ANOVA, Interaction: F _2, 11_ = 15.16, p < 0.0007, Group: F _2, 11_ = 15.16, p < 0.0007; Time: F _1, 11_ = 2.17, p = 0.17, control vs. ChR2, p < 0.05; control vs. GtACR2, p < 0.001,). Error bar indicates ± s.e.m. *** P < 0.001. *p < 0.05.

### Working model

Both SPNs and FSIs are known to receive convergent inputs from different cortical regions ^31, 32^. We hypothesize that these inputs can provide information on distance to target. Based on the anatomy of the FSI-SPN circuit and our results, we propose a working model for how this circuit contributes to pursuit behavior (**Fig. 7)**. According to this model, the SPNs and FSIs that form a functional unit share excitatory cortical inputs. Signals representing target distance are sent via the corticostriatal projections to FSIs and SPNs. FSI output sends a slightly delayed of this signal to the relevant SPNs. Consequently, a given velocity SPN can receive two inputs, one net positive signal from the cortex that represents distance, and a slightly delayed version of the same input from the FSI. Because the FSI-SPN projection is inhibitory (**Extended Data Fig. 11**), it is subtracted from the excitatory input, and the SPN output will then reflect the difference between excitatory cortical input and a temporally delayed version of the same signal. This ‘temporal difference’ signal, similar to taking the time derivative of the distance representation, is then used as a velocity command to drive performance. At rest velocity is close to zero, consistent with the typically low baseline firing rates of SPNs ^33^. By contrast, FSIs are characterized by high tonic firing rates, which make them more suitable for signaling the distance variable. We found that tonic FSI firing close to the middle of the range is used to represent the effective zero, when the target is directly in front of the mouse. Different types of errors in distance control (left vs. right) are indicated by the sign of the signal relative to the effective zero (**Fig. 3**) ^34, 35^. For neurons that increase firing when the target is to the left, the left extreme is represented by the highest firing rate and the right extreme is represented by the lowest firing rate, and vice versa for neurons that increase firing when target is on the right side. A positive error means that distance error has increased, i.e. the target is farther away in a particular direction, and velocity in that direction must be increased accordingly to reduce the distance error. Thus an increase in left distance increases the temporal difference signal computed by the FSI-SPN circuit, and a corresponding increase in the leftward velocity command. A negative difference signal indicates that the distance error is decreasing and velocity in that direction should be reduced. SPN output is therefore regulated by the *change* in distance between self and target in real time.

**Fig. 7.**
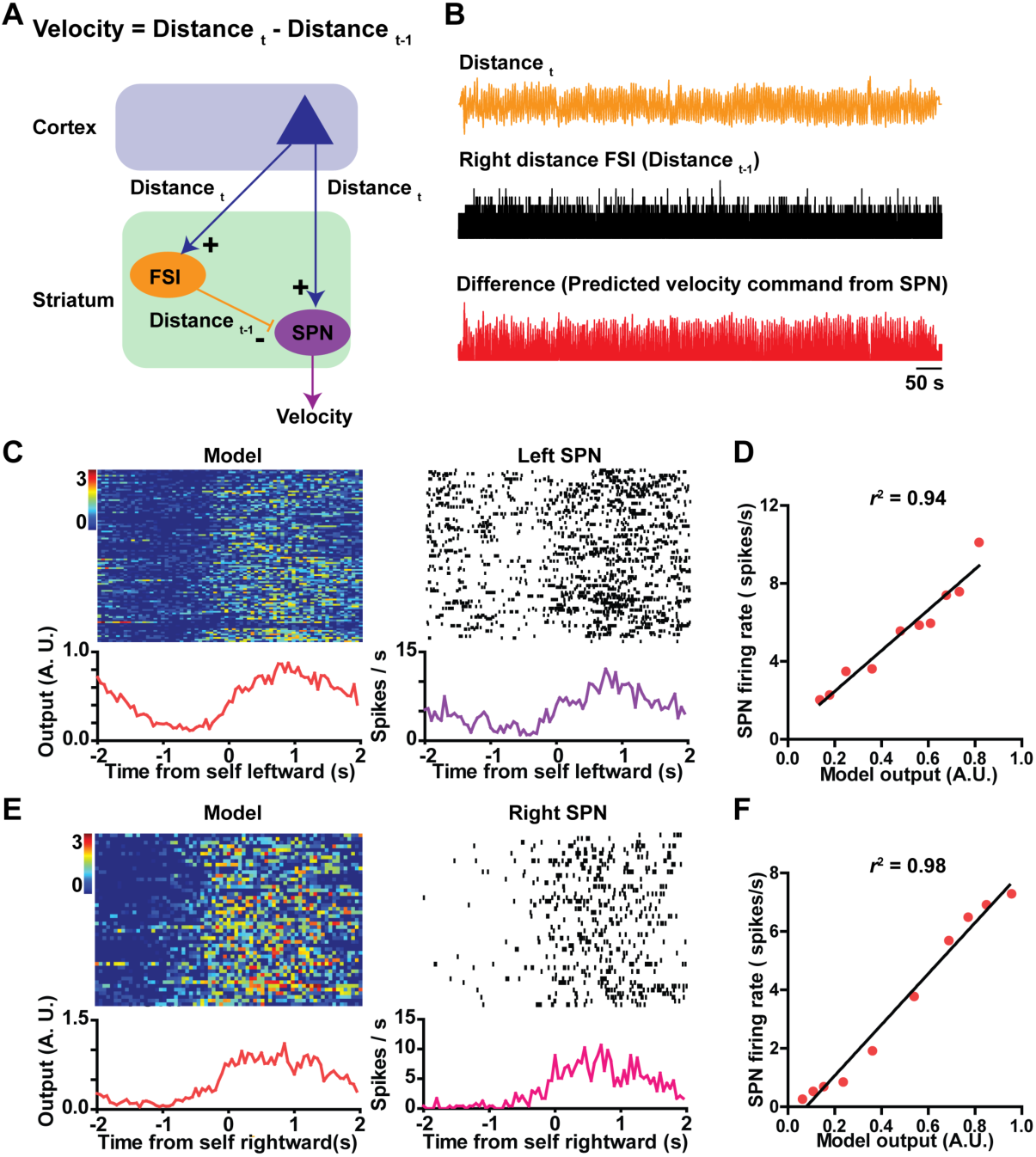
Working model of how FSI-SPN circuit contributes to continuous pursuit. a) Schematic diagram of the FSI-SPN feedforward inhibition circuit. Note that in the equation for velocity, distance refers to distance to target. This equation only applies to self-velocity during continuous pursuit. As shown here, excitatory cortical inputs reach both the FSI and SPN. In this case, some excitatory input represents distance to target, a key error signal used to guide pursuit behavior, and the FSI relays a slightly delayed version of the same signal to the SPN. Thus the delayed distance signal (Distance _t-1_) is subtracted from the distance signal (Distance _t_), and the SPN output reflects the difference between these two signals, i.e. the change in distance in this time step. As the feedforward inhibition circuit can function as a differentiator, the velocity command is proportional to the change in distance in a direction-specific manner. b) Actual traces from our experiments illustrating the variables used in the model. c) The model is compared with actual data using simultaneously recorded left distance FSI and left velocity SPN from the same mouse (same hemisphere). Using actual distance measure as an estimate of the Distance _t_ signal (shifted by 100 ms to account for the delay in the perceptual system) sent to the striatum, and FSI activity as the measure of the Distance _t-1_ signal sent to the SPN, a subtraction generates a difference signal (model) that is compared to the activity of an actual left velocity SPN. d) Model output is highly similar to right velocity SPN output (Pearson’s r, p <0.0001. e) Same as c, except that the model is generated using a right distance FSI. The model is compared with actual data using simultaneously recorded right distance FSI and right velocity SPN. f) High correlation between model output and right velocity SPN output (p <0.0001).

Our working model is supported by optogenetic manipulation of SPNs. We used SIO-GtACR to inhibit SPNs in the direct (D1+) and indirect (A2A+) pathways (**Extended Data Fig. 12**). Inhibition of SPNs in either pathway did not significantly change distance to target. However, direct pathway inhibition reduced velocity and increased distance error, whereas indirect pathway inhibition slightly increased velocity.

## Discussion

To successfully pursue a moving target, it is necessary not only to regulate velocity and direction of movement but also to continuously monitor the distance between self and target. Here we showed for the first time that mice use internal representations of distance to guide their pursuit behavior (**Fig. 1**). We also found that a striatal microcircuit is critical for pursuit behavior. Striatal FSIs not only represent target distance, and are also necessary for successful pursuit performance. Their projections to the downstream SPNs are critical in regulating self-velocity during pursuit.

In our task, the target always moves from side to side, each cycle consisting of both leftward and rightward movements. At any time, the target could be on either side of the mouse. We identified distinct types of self-target distance representations in FSIs. Some neurons increase firing when the target moves to the left side of the animal, and decrease firing when it moves to the right, and vice versa for other neurons (**Fig. 3**). On the other hand, SPNs are strongly correlated with movement velocity, with clear direction-specificity (**Fig. 2**).

In order to test whether FSIs are necessary for pursuit, we used a variety of methods to manipulate their activity. Silencing these neurons using hM4Di or TeLC resulted in impaired pursuit (**Fig. 5**). However, as these techniques lacked temporal precision, it was only possible to measure overall distance and velocity during the session. It was not possible assess the immediate impact of FSI silencing on pursuit performance. Using optogenetics, we were able to obtain a better understanding of how bidirectional manipulation of FSI activity affected pursuit behavior. In particular, inhibition using GtACR2 reduced distance while excitation using ChR2 increased distance. As expected, these manipulations have the opposite effects on velocity given the mainly inhibitory effects of FSIs on SPNs (**Fig. 6**). It should be noted that, while these effects support the hypothesis that FSIs are critical for pursuit, it is unclear whether sensory detection in general was affected by manipulation of FSI activity. That is, we cannot easily rule out the possibility that FSIs are also critical for distance perception per se. The role of striatal FSIs in sensory processing remains to be elucidated.

Interestingly, recent studies have also elucidated neural substrates underlying natural pursuit behavior. A key area that has been reported to be important for pursuit of prey in hunting behavior is the periaqueductal gray (PAG). This area is the target of descending projections from many regions, and recent work has shown that projections from the central amygdala and hypothalamus can activate pursuit behavior^36–38^. Interestingly, PAG also receives inputs from the BG ^39^. It is therefore possible that the BG provide another source of descending projections to the PAG that can also modulate pursuit behavior.

Our results suggest that tonic FSI firing close to the middle of the dynamic range is used to represent the effective zero in horizontal distance, when the target is directly in front of the mouse. Different types of errors (left vs. right) are indicated by the sign of the signal in relation to the effective zero. For example, for neurons that increase firing when the target is to the left, the left extreme corresponds to the highest firing rate and the right extreme to the lowest firing rate, and vice versa for neurons that increase firing when the target is on the right side. These two types of FSIs may project to distinct types of SPNs that signal velocity. A distance error signaling ‘too much to the left’ would suppress rightward neurons and activate leftward SPNs, whereas ‘too much to the right’ would suppress leftward neurons and activate rightward SPNs. This organization would enable bidirectional control by FSIs.

In our working model, the feedforward FSI circuit allows differentiation of the distance variable during tracking, converting it into instantaneous velocity commands from SPNs. Previous optogenetic experiments showed that movement velocity depended on the frequency of striatonigral stimulation ^19^. We also showed that inhibition of the direct pathway (D1+) neurons significantly reduced velocity during pursuit, but inhibition of indirect pathway (A2A+) neurons did not. Thus the distance to velocity conversion is achieved mainly by the direct pathway. Previous work also showed that GABAergic output from the substantia nigra pars reticulata represents instantaneous position vector components ^34^. Using the direct pathway, velocity commands could be integrated into reference signals representing proprioceptive position references that change over time ^34, 35, 40–43^. These reference signals from the BG output would then reach downstream proprioceptive position control systems for posture and body configuration ^44, 45^.

Although velocity and distance are both continuous, time-varying variables, there are key differences between them. Movement velocity, detectable by mainly proprioceptive and vestibular sensory inputs ^40^, is close to zero at rest. The usually low firing rates of SPNs are ideal for representing this type of variable ^33^. On the other hand, distance, usually detected by teloreceptive inputs, is rarely close to zero. FSIs are characterized by high tonic firing rates, which make them ideal for signaling distance. Presumably inputs from sensory cortical areas provide distance information that is sent to the striatal FSIs. Visual inputs are obviously critical, but we cannot rule out other sensory modalities. Based on our results, we hypothesize that representations of specific pursuit errors can activate the appropriate combination of direction-specific SPNs (e.g. moving leftward when the target is on the left side) and suppress the conflicting SPNs. With their strong inhibitory projections to SPNs, FSIs can suppress the activation of irrelevant or antagonistic SPNs via monosynaptic inhibition. On the other hand, via disinhibition or perhaps excitatory effects of GABA ^46^, they can also have net excitatory effects on downstream SPNs.

Given the importance of approaching a spatially discrete target in normal behavior, our results suggest how striatal FSIs have been implicated in so many functions ^14–16^. For example, although reward is the eventual goal of behavior in our task, the striatal neurons we recorded represented specific spatial variables necessary for successful performance, rather than reward prediction or consumption. In accord with previous work ^17^, we found that, in the sensorimotor striatum at least, FSI and SPN activity is largely independent of reward delivery. Although some studies have suggested that striatal neurons are modulated by reward, they did not monitor continuous behavioral variables^47^. It is impossible to explain our results in terms of predicted reward value because FSI activity reflects distinct distance errors that are spatially defined, e.g. left and right. Caution is needed in interpreting experiments that lack precise and continuous behavioral measures. Abstract psychological concepts like reward value must be well anchored to clear experimental measures to generate interpretable experimental results.

Our results also resolve current controversies on the relationship between striatal activity and behavior. A recent study by Owen et al concluded that FSIs are not needed for behavioral performance but for learning and plasticity ^48^. But that study did not use continuous behavioral measures and consequently failed to detect the crucial relationship between FSI activity and key performance variables like distance to target. In fact, because their learning tasks involved orienting and approaching reward targets, any learning defects observed following disruption of FSI output can be explained by performance deficits in approaching the target. Moreover, while some recent studies observed correlation between velocity and striatal output ^18, 49^, this conclusion has also been questioned. For example, Klaus et al found that SPNs can encode action identity independent of velocity ^50^, but their conclusion is not supported by the data, which consisted of accelerometer measures with little spatial information and calcium imaging with low temporal resolution, which cannot reveal the relationships between behavior and neural activity. Moreover, free behavior in an open field is exceedingly complex and not always dependent on striatal output, so caution is needed in interpreting neural activity recorded during such behavior.

On the other hand, while classic studies discovered SPNs linked to movement speed ^51 52^, they did not measure behavior continuously in freely moving animals and relied on average measures. In fact, speed is not the appropriate measure, though it is correlated with neural activity. Kim et al were the first to show that SPNs can represent vector components of velocity in freely moving animals, and to demonstrate that such representations are similar whether or not the behavioral outcome is rewarding or aversive^17^. Just like a recent study using calcium imaging in head restrained mice ^53^, Kim et al concluded that both SPNs and FSIs represented velocity. However, the video-based motion tracking used in their study did not have sufficient spatial and temporal resolution to dissociate the different contributions of SPNs and FSIs. In the present study the use of 3D motion capture reveals that, during tracking, FSIs more commonly represent distance to target, while many sensorimotor SPNs represent velocity.

The velocity representation in a large proportion of sensorimotor SPNs is therefore unique, and so far not found in any other cell type except nigrostriatal dopamine neurons^41^. Striatal velocity neurons have strong preference for direction of egocentric motion (left, right, up, down). It is important to note that conventional notions of speed or vigor cannot begin to capture the richness of kinematic representations in SPNs. Actual volitional movement is shaped by a combination of distinct vector components (at least 4 major classes depending on the direction of movement). A population of dif ferent types of velocity-related neurons can therefore represent a velocity vector.

In addition, although previous work found that SPNs are typically selective for contraversive movements^54^, we did not find more contraversive neurons than ipsiversive neurons in our study. There are two possible explanations for this discrepancy. First, we might not have sampled enough neurons to obtain an accurate estimate of the distribution of contraversive and ipsiversive neurons. Secondly, the velocity-related SPNs recorded in this study might not be the same as those involved in spontaneous behavior. While they could overlap, additional neurons that receive bilateral projections from the cortex may be recruited for the pursuit task.

A striking feature of our results is the high correlations between single unit activity and behavioral variables. High correlation is crucial if a signal like firing rate is used in analog computing, as is the case in our model. The concept of correlation is often misunderstood in contemporary neuroscience, in part because traditional studies have consistently failed to find high correlations between neural activity and behavior. Low correlation in most studies are responsible for the development of many elaborate statistical techniques and for conflicting interpretations about the neural code ^55^. For example, multiple regression is often used to analyze single unit data, and many argue that neural signals multiplex of multiple behavioral variables, without explaining how to demultiplex the signals. In practice this allows one to argue that any neural signal can multiplex just about any collection of variables. Such a conclusion is hardly falsifiable. It is important to emphasize that, in order to observe the relationship between neural activity and behavior as reported here, it is critical to quantify behavior accurately and continuously in unrestrained animals, and to use measures with high temporal and spatial resolution. Our results suggest that low correlation between neural activity and behavior is due at least in part to limitations in behavioral measures, the widespread use of head restraint, and conventional experimental designs that rely exclusively on categorical classification of behaviors. In contrast, our working model, which involves simple analog computing, has the advantage of being parsimonious and falsifiable.

In conclusion, while our results do not exclude other functional roles for striatal output^56^, they do demonstrate the role of sensorimotor SPNs in the control of velocity during continuous pursuit, and show that the FSI-SPN circuit implements the distance-velocity transformation, the key computation required for the precise guidance of volitional behavior.

## METHODS

### Animals

All experimental procedures were approved by the Animal Care and Use Committee at Duke University. Male and female C57BL/6J mice were used in all experiments. For TeLC and optogenetic experiments, Parvalbumin-Cre mice (Pvalb-2A-Cre-D) and D1-cre (Drd1a-Cre) and A2A-cre (Adora2a-Cre) mice were used. For DREADD experiments, Drd1a-tdTomato:: *PV*-Cre mice were used. Drd1a-tdTomato reporter mice were generated in the Calakos lab (RRID: IMSR_JAX:016204)^57^. All mice were between 3-8 months old. They were maintained on a 12:12 light cycle and tested during the light phase. The mice used for electrophysiology experiments were singly housed; others were group housed. During all behavioral experiments, the mice had restricted access to water and food. After each training and recording session, they had free access to water for ∼ 30 minutes and ∼3 g of home chow. Their weights were maintained at 85-90% of their *ad libitum* weights.

### Behavior task and analysis

The moving reward target is a sucrose spout moved by a stepper motor from left to right relative to the mouse standing on a platform (Bipolar, 56.3 x 56.3 mm, DC 1.4A, 2.9Ω, 1.8 degree / step, Oriental motor, USA). The stepper motor was controlled using MATLAB (Mathworks). Two infrared reflective markers (6.35 mm diameter) were used for tracking mouse and target movements. One marker was located on the wireless recording headstage and another was approximately 20 mm from the sucrose spout. These markers were captured by eight Raptor-H Digital Cameras (Motion analysis, CA, 100 Hz sampling rate). The data was transformed into Cartesian coordinates (x, y and z) by the Cortex program (Motion Analysis, CA). MATLAB communicated with the Cortex program (Motion Analysis) online to control reward delivery based on mouse behavior. Reward (∼12 ul of 20% sucrose) was delivered every 800 ms during following. Prior to the recording sessions, all mice were trained until they followed the target consistently (2-4 hours of training over 1 week). In most experiments, the target moved at a constant velocity (16 mm/s). On some sessions, however, target velocity was varied randomly (5-48 mm/s, updated every 2 ms by MATLAB).

### Wireless *in vivo* electrophysiology

A total of 24 C57BL/6J mice were used in the electrophysiology experiments (17 males and 7 females). We used custom-built 16-channel microwire arrays with micro-polished tungsten wires, 35 µm in diameter and 4-5 mm in length, in a 2 by 8 configuration, and attached to an Omnetics connector (Omnetics Connector Corporation) as well as arrays with a similar configuration from Innovative Neurophysiology. To implant the electrode arrays, each mouse was anesthetized with isoflurane (induction at 3%, maintained at 1%), and head-fixed on a stereotax (Kopf). Meloxicam (2 mg/kg) was administered subcutaneously after anesthesia induction and prior to surgery for pain relief. Detailed surgery procedure was previously described ^43, 58^. The stereotaxic coordinates are (mm relative to Bregma): AP +0.4 mm, ML ±2.4 mm, DV −2.3 mm. Following the surgeries, all mice were allowed to recover for at least two weeks before recording neural activity. Single-unit activity was recorded with a miniaturized wireless headstage (Triangle Biosystems) and a Cerebus data acquisition system (Blackrock), as described previously ^59^. The data was processed using online sorting algorithms and then re-sorted offline (Offline Sorter, Plexon). When classifying single unit waveforms, the following criteria were used: 1) a signal to noise ratio of at least 3:1; 2) consistent waveforms throughout the recording session; 3) refractory period of at least 800 µs. For correlation analysis, we first obtained rate histogram data in 10 ms bins. For FSIs, a 200 ms Gaussian filter was used for smoothing. For SPNs, a 50 ms Gaussian filter was used for smoothing. Correlation with each of the different behaviorally relevant variables (distance, velocity, and acceleration, and reward delivery) was computed. For reward correlation, neural and reward times were aligned by rightward self movement from –3.5 s (left) to 3.5 s (right). Using different peri-event windows does not change the results. We compared the automatically computed correlation coefficients for different behavioral variables. A neuron is considered to be correlated with a particular behavioral variable if the absolute value of r is higher than 0.75 and higher than the r values for alternative variables.

### *In vivo* calcium imaging

Five PV-Cre mice were used for calcium imaging experiments. Each mouse was unilaterally injected with 500 nl of AAV-EF1a-DIO-GCamp6s (Stanford virus core) into dorsolateral striatum (Bregma +0.4 mm, ±2.4 mm; −2.3 mm) every 0.2 mm from 3.0 mm using a Nanoject III injector (Drummond Scientific, USA) at a rate of 1 nl per second. The injection pipette was left in place for 10 min post-injection before it was retracted. After viral injection, a gradient index (GRIN) lens (Inscopix: 1 mm x 9 mm, n = 3; UCLA Miniscope, 1.8 mm x 4.3 mm, n =2) was implanted in the dorsolateral striatum directly above the injection site after aspirating ∼2 mm of the above cortical tissue with 21-gauge blunt needle. The lens was then secured to the skull using dental cement and covered with Kwik-Sil to protect the lens surface. Two weeks after the GRIN lens implantation, the baseplate (Inscopix or UCLA Miniscope) was mounted onto the mouse head under visual guidance to determine the best field of view. All recorded movies of calcium activity were initially preprocessed in Mosaic (Inscopix) for motion correction and spatial binning (5 x 5 for Inscopix; 4 x 4 for miniscope), and subsequently analyzed using custom MATLAB scripts. We used constrained non-negative matrix factorization (CNMF) for denoising, deconvolving, and demixing of calcium imaging data ^60, 61^. This method allows accurate source extraction of cellular signals.

### TeLC experiments

Cre-inducible adeno-associated viral (AAV) plasmids containing GFP-tagged TeLC or GFP alone, each with the reading frame inverted in a flip-excision (FLEX) cassette (AAV-FLEX-TeLC and AAV-FLEX-GFP) were originally obtained from Dr. Peer Wulff, Christian Albrechts University, Kiel, Germany ^25^, and the AAV vector was generated by the University of Pennsylvania vector core facility in serotype AAV2/5. We first implanted cannulae in PV-Cre mice (n = 13, 24 gauge, 3-mm length below pedestal, Plastics One; AP +0.4 mm, ML ±2.4 mm, DV −2.3 mm). Cannulae were secured in place with skull screws and dental acrylic. A stylet was inserted and protruded ∼0.2 mm beyond the end of each cannula. The mice then trained daily for ∼10 days, each session consisting of 200 side to side movements of the target. To inject the virus (0.5ul of either TeLC-GFP or GFP virus), the stylets were removed, and virus was infused through custom made 31 gauge steel injectors extending ∼0.5 beyond the tip of the guide cannulae at 0.1ul/min for 5 min. The injector was kept in the brain for at least 5 min for diffusion. Coordinates for all injections relative to bregma were as follows: AP + 0.4 mm, ML ±2.4 mm, DV −2.3 mm ^62^. Approximately 3 weeks after surgery, the mice were tested on the pursuit task again.

### DREADD experiments

The hM4Di virus (n = 6) or EYFP virus (n = 6) injected into the sensorimotor striatum of *Drd1a*-tdTomato::*PV*-Cre mice. The *CAG*-FLEX-*rev-*hM4D:2a:GFP plasmid was provided by Scott Sternson (Addgene #52536). UNC Viral Vector Core packaged this plasmid into AAV 2/5 and also provided AAV2/5-EF1a-DIO-EYFP (titers > 1 x 10^12^ particles/mL). Small craniotomies were made over the injection sites and 1.0 µL of virus was delivered bilaterally to dorsolateral striatum via a Nanoject II (Drummond Scientific) at a rate of 0.1 µL/min. The injection pipette was held in place for 5 minutes following injection and then slowly removed. Coordinates for all injections relative to bregma were as follows: AP: +0.8 mm, ML: ± 2.7 mm, DV: −3.2 mm. Mice were allowed a minimum of 3 weeks before behavioral training. The behavioral procedure is the same as described above, except mice were first tested on 100 cycles of target motion. This pre-injection period establishes the baseline performance measures for each mouse, before given injection of the synthetic ligand clozapine-N-oxide (CNO) for the hM4Di receptor. CNO was dissolved to 10 mg/mL in DMSO and diluted in PBS solution to administer 5 mg/kg per mouse with a maximum injection volume of 0.5 mL. The same volume of PBS/DMSO solution was used as the control vehicle injection. 30 minutes after control or CNO injection, the mice were tested again for 100 cycles.

### Optogenetics

To determine the effect of PV manipulation, fourteen Pv-Cre mice were used in the optogenetic experiments (PV-Cre control, n = 4; ChR2, n = 4; stGtACR2, n = 6). In addition, to determine the effect of SPN manipulation, ten mice were used in the optogenetic experiments (D1-cre (stGtACR2), n = 4; D2-Cre (stGtACR2), n = 3; Control, n =3). Adeno-associated viral vectors were used for Cre-dependent expression of the excitatory channelrhodopsin (pAAV-EFIa-DIO-hChR2(H134R)-EYFP, Duke viral core, titer > 1 x 10^12^ particles/mL), the inhibitory channelrhodopsin stGtACR2 (soma-targeted Guillardia theta anion-conducting channelrhodopsin 2, pAAV_hSyn1-SIO-stGtACR2-FusionRed, titer > 1 x 10^13^ particles/mL, Addgene). The vectors were injected (0.5 µl each side) using the same procedure as described above using the following coordinates: AP + 0.4 mm, ML ±2.4 mm, DV −2.8 mm. Both heterozygous male and female PV-Cre mice were used. Photo-stimulation is always bilateral. To measure *in vivo* neural activity during photo-stimulation, we implanted an optic fiber just above the site of injection (DV −2.3 mm) on one side and an optrode on the other side. The optrode was a custom made microwire array (4-mm, 4 x 4 tungsten) with an optic fiber attached to the lateral side and angled so the cone of the emitted light encompassed the electrode tips ^63^. Control mice received bilateral implants of optic fibers only. Before testing each day, mice were connected to a 2-m sheathed fiber (105-µm core diameter, 0.22 NA) with a ceramic sleeve (Precision Fiber Products). The fiber was connected to a 470 nm DPSS laser (Shanghai Laser & Optics), controlled with MATLAB (MathWorks). Before the experimental session, the final output of the laser was adjusted, based on the transmittance of each implant, to be 8-10 mW into the brain. Photo-stimulation (500 ms) was delivered 3.2 s after at least 800 ms of following. If the mouse continues to follow, stimulation was delivered every 3.2 s. For optrode recordings, tethered setup was used (Cerebus data acquisition system, Blackrock).

### Histology

After the completion of all behavioral tests, the mice with chronically implanted electrode arrays were deeply anesthetized with isoflurane and transcardially perfused with 0.9% saline followed by 10% buffered formalin solution. Coronal brain sections (80 µm) were sliced using a Vibratome 1000 Plus, stained with thionin, and examined with a light microscope to verify location of the electrode tips within the striatum. Mice were deeply anesthetized with isoflurane, perfused transcardially with Tris buffered saline (TBS; pH 7.4) containing 25 U/ml heparin, followed by 4% paraformaldehyde (PFA) in TBS. Perfused brains were removed, post-fixed overnight at 4°C in 4% PFA, and then cryo-protected with 30% sucrose in TBS for 48 hrs. Brains were cut into 50 µm coronal sections using a cryostat (Leica CM 3000), and treated with blocking solution (TBS containing 5% normal goat serum and 0.2% Triton X-100) for 2 hr and incubated overnight at 4°C with chicken anti-GFP monoclonal antibody (1:500; abcam; ab13970). After washing three times with TBST (TBS containing 0.2% Triton X-100), sections were incubated with goat anti-chicken Alexa Fluor® 488 IgG (1:500; invitrogen; A11039) overnight at 4°C. Sections were counterstained with a 4’, 6-diamidino-2-phenylindole solution (DAPI; Sigma-Aldrich). After washing four times, the sections were cover slipped with FluorSave (CalBioChem) aqueous mounting medium. Whole brain section images were taken by tile scan imaging using a LSM 710 confocal microscope (Zeiss) with a 10x objective. To observe individual neurons expressing ChR2, stGtACR, TelC, or DREADD, a 20x objective lens with 3× digital zoom was used.

### Statistics

Statistical analysis was performed using Matlab and Graphpad Prism. No statistical methods were used to predetermine sample size. All statistical tests were two-tailed. To determine the effects of FSI inactivation, we used two-way repeated measures ANOVAs. Post hoc tests were performed whenever there was a significant interaction. To compute correlation between neural activity and behavioral variables, we used data from the entire session. First, we divided the data based on whether the animal was following the target, as described above. We then computed the task relevant behavioral variables, which include distance, velocity, and acceleration. We sorted the array of both firing rate and kinematics and excluded any outliers at the two extremes (<1% or >99%). We then divided the values into 20 bins and calculated correlation coefficient (Pearson’s r) between the kinematics and neural activity using mean values from the bins. For the calcium imaging experiment, data was divided into 6 bins. For correlation with reward (sucrose drops/s), we calculated correlation coefficient only when animals were following because the reward was not delivered if they did not follow the target.

### Data availability

All data and Matlab codes used in the present study are available upon request.

## Supporting information

Video 1 following

Video 2 not following

Video 3 calcium imaging

## Acknowledgements

We would like to thank Justin O’Hare and Nicole Calakos for their help with DREADD experiments, Koji Toda and Dongye Lu for help with 3D printing, Scott Soderling for providing the Inscopix microscope for some of the calcium imaging experiments, and Mark Rossi and Konstantin Bakhurin for helpful comments on the manuscript. This research was supported by grants from the National Institutes of Health (DA040701, NS094754, MH112883 to HHY, DA033610 to AEW, and MH117429 to IK). The authors declare that there is no conflict of interest regarding the publication of this article.

## Author Contributions

NK and HHY designed the experiments. DG and AW provided reagents. NK, HEL, RH, GW, and IK conducted experiments. NK and HHY analyzed the data and wrote the manuscript.

---

**Video 1. Representative 3D motion capture of a mouse during pursuit behavior (following).**

**Video 2. Representative 3D motion capture of a mouse that is not following the target (same mouse as in Video 1).**

**Video 3. Representative calcium imaging signals using the UCLA Miniscope.**

## Extended Data Figures

**Extended Data Fig. 1.**
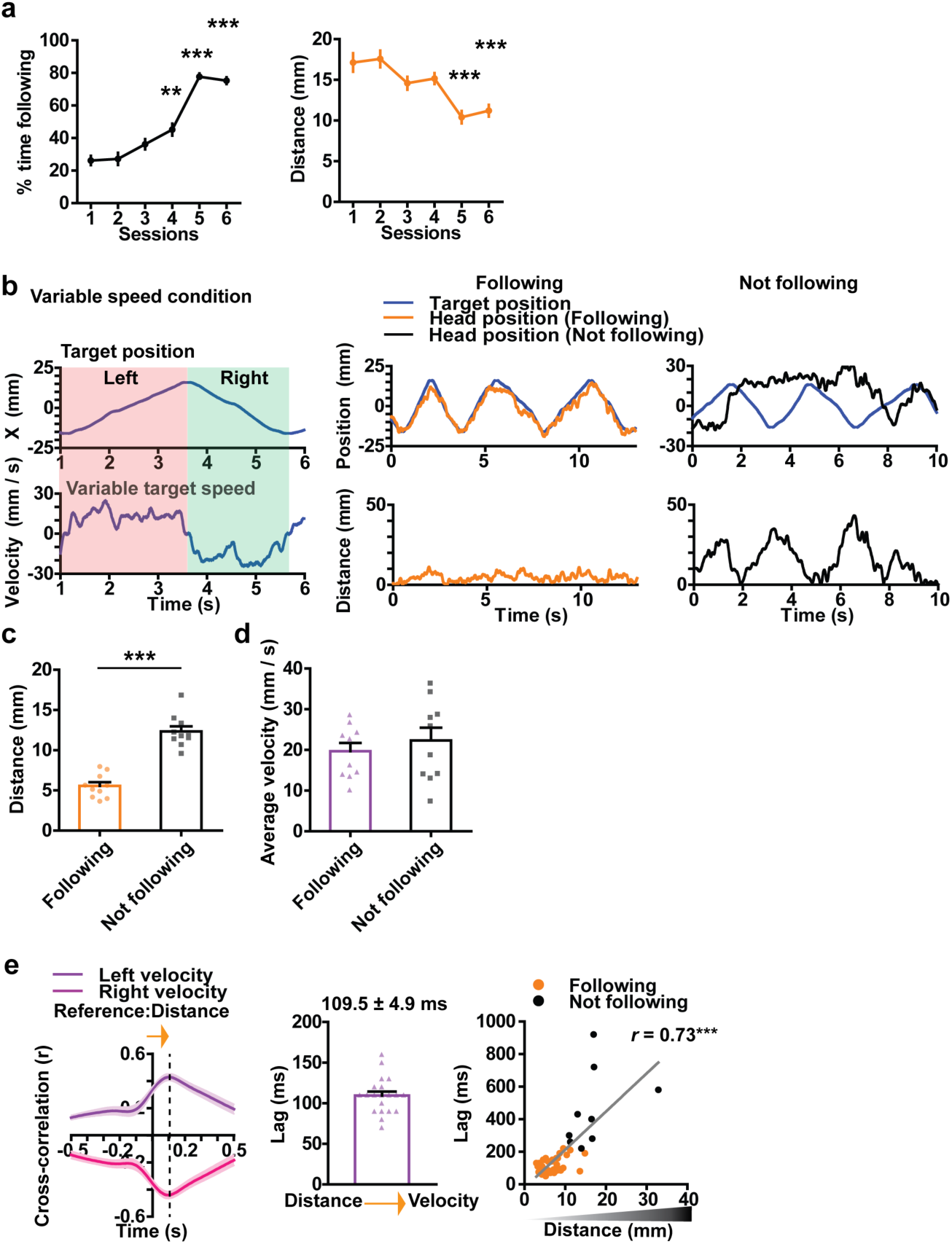
Behavior summary. a) In 6 mice, we recorded behavior with 3D motion capture during acquisition of the continuous pursuit task. Behavioral performance improved in the course of training. Time spent following the target increased significantly (Repeated measures ANOVA, main effect of sessions, F _5, 35_ = 59.10, p < 0.0001). Distance error is reduced in the same period (Repeated measures ANOVA, main effect of sessions, F _5, 35_ = 16.76, p < 0.0001). b) Representative traces of mouse head and target position during a session with variable target velocity. Left panel shows the head and target positions during ‘Following.’ Right panel shows the head and target positions during ‘Not following’ in which two positions are different. c) Self-target distance error is significantly lower during following than not following (paired t-test, p < 0.0001). Error bars indicate ± s.e.m. *** p < 0.0001. d) Average velocity is similar between following and not following (p = 0.053). e) Illustration of the temporal relationship between distance and self-velocity. Cross correlation analysis reveals the lag between these two variables, showing that distance leads self-velocity during pursuit behavior. In addition, the lag shows a positive correlation with distance (r = 0.73, p < 0.0001). When the mouse is actively pursuing and maintaining a small distance, the lag between distance and velocity is small, but the lag increases as pursuit performance declines. Dashed line indicates the time of peak positive and negative correlation. Shaded areas indicate ± s.e.m.

**Extended Data Fig. 2.**
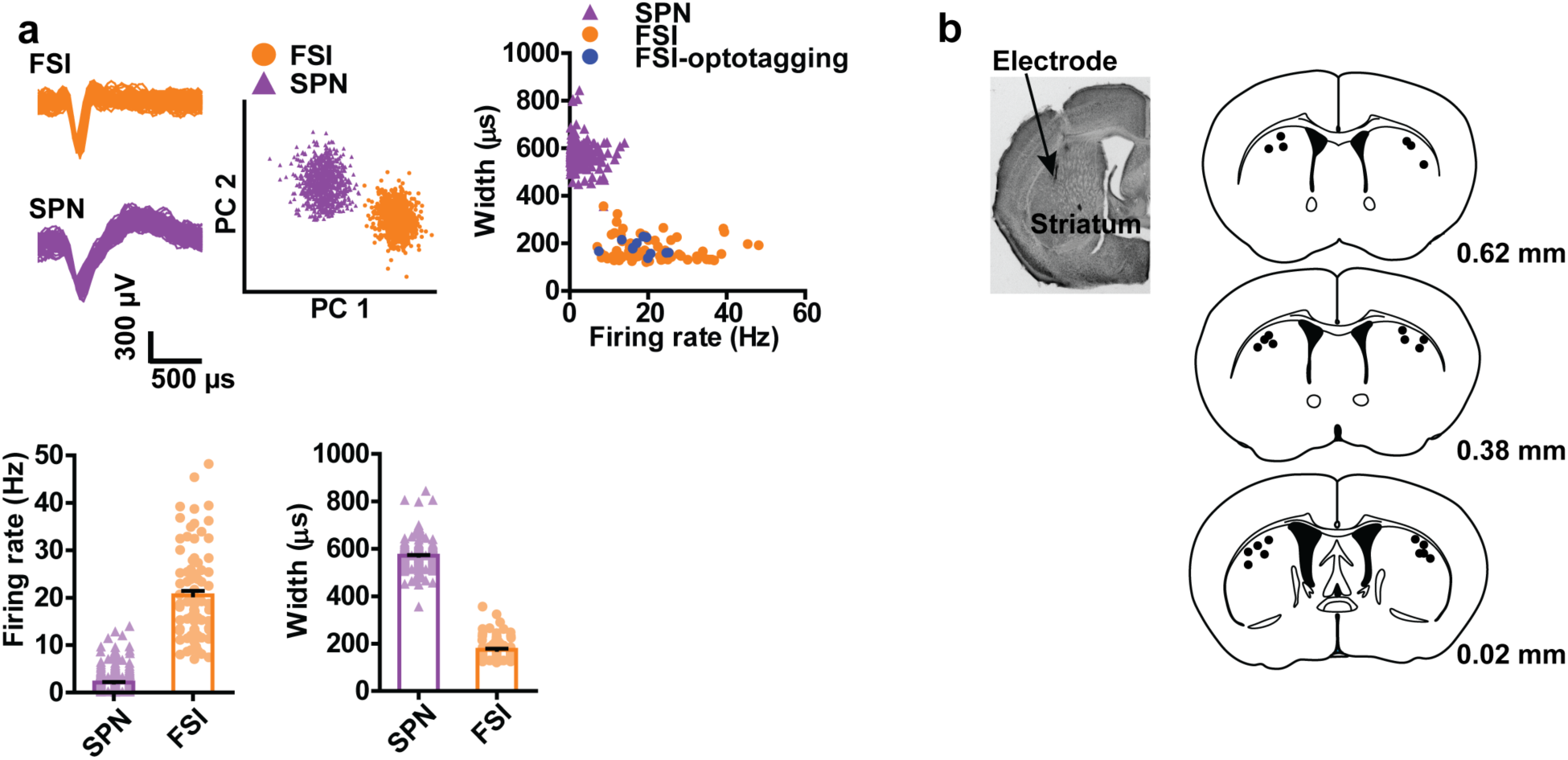
Summary of in vivo electrophysiology. a) SPNs and FSIs are classified based on their spike waveforms and firing rates. b) Placement of electrodes in the sensorimotor striatum.

**Extended Data Fig. 3.**
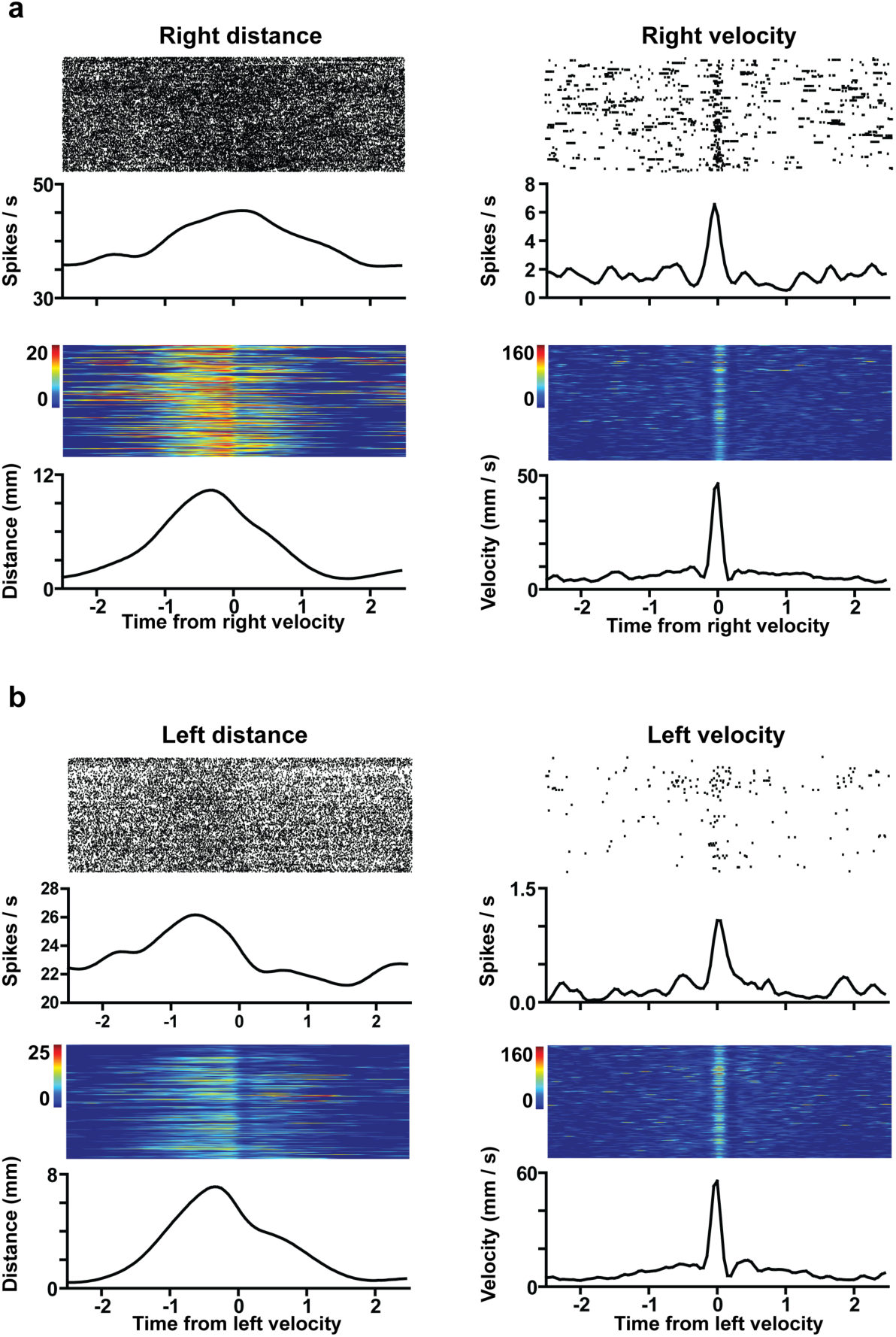
Distance and velocity classification. a) A representative examples of right distance FSI and right velocity SPN from the same animal aligned by right velocity. It is clear from these examples that, while distance and velocity co-vary, these two variables can also be dissociated. FSIs, which show typical tonic activity, are suitable for representing distance, whereas SPNs, which show sparse firing, are suitable for representing velocity. b) A representative examples of left distance FSI and left velocity SPN from the same animal aligned by left velocity.

**Extended Data Fig. 4.**
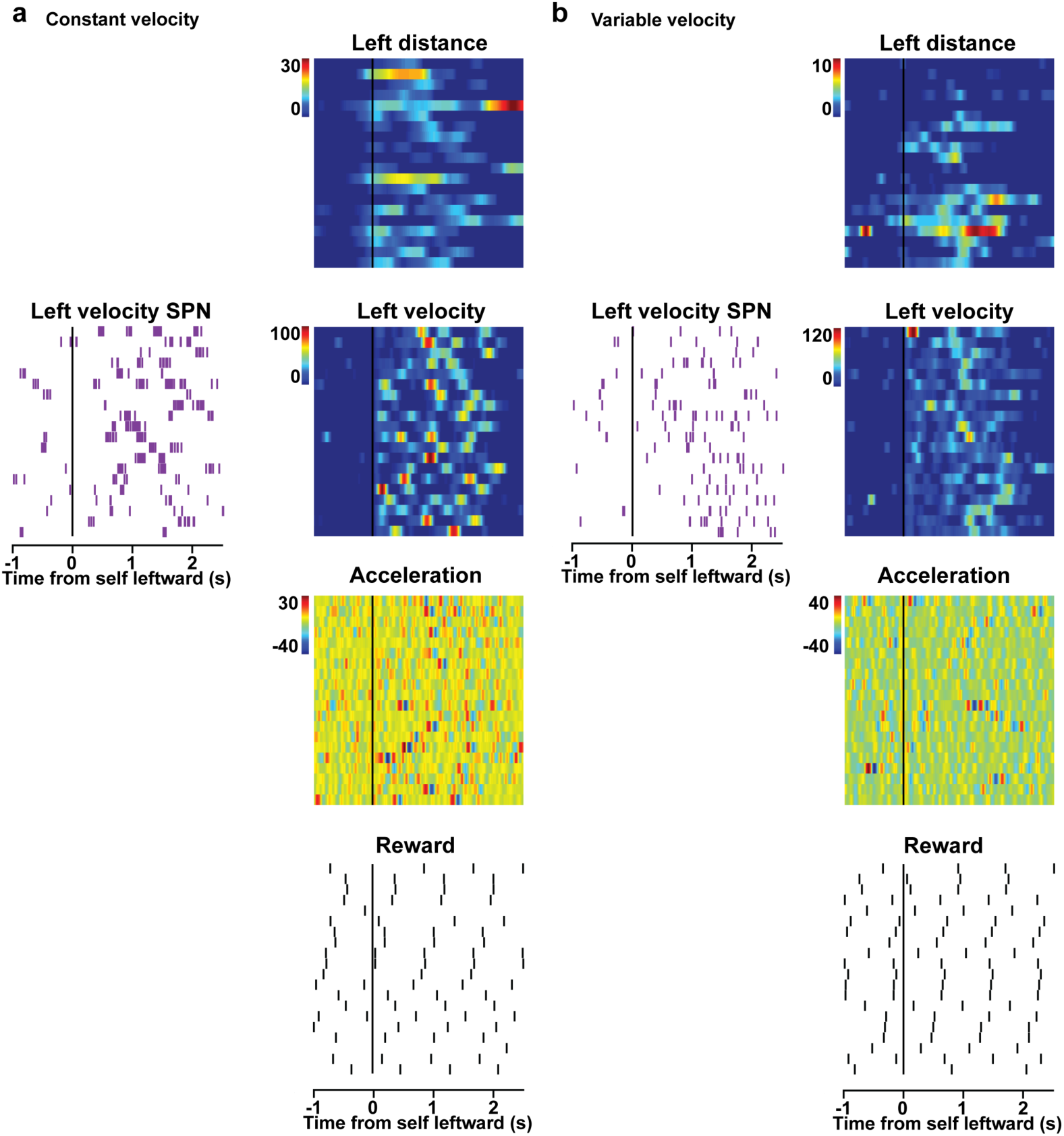
Representative velocity-correlated SPN. a) In constant target velocity condition, correlation between a representative left velocity SPN and different behavioral variables (distance in mm, velocity in mm/s, acceleration in mm/s^2^, and reward in sucrose drops, 20 consecutive target movements while the animal is following). b) In variable target velocity condition, correlation between a representative left velocity SPN and different behavioral variables.

**Extended Data Fig. 5.**
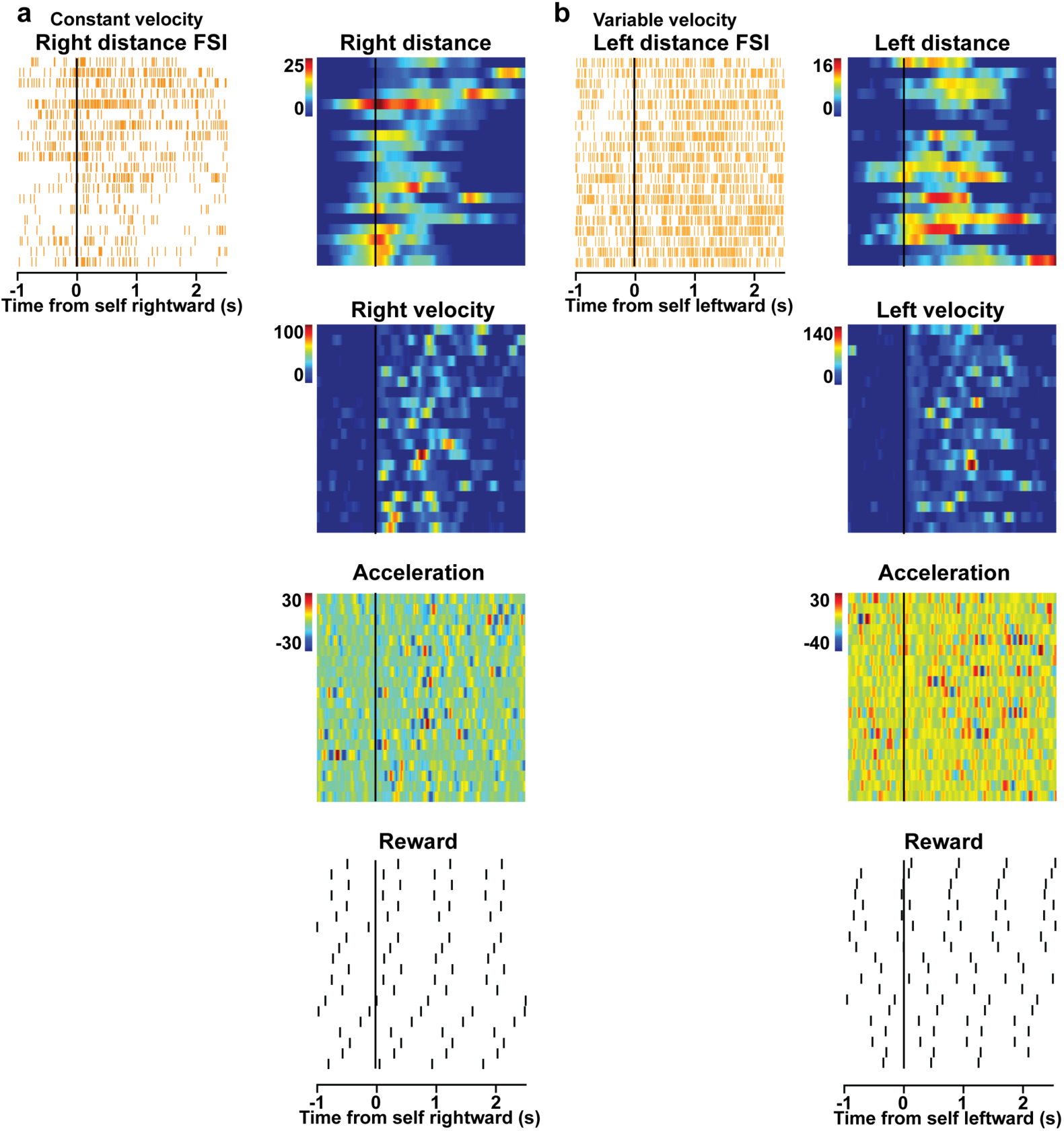
Representative distance-correlated FSI. a) In constant target velocity condition, correlation between a representative right distance FSI and different behavioral variables ((distance in mm, velocity in mm/s, acceleration in mm/s^2^, and reward in sucrose drops, 20 consecutive target movements while the animal is following). b) In variable target velocity condition, correlation between a representative Left distance FSI and different behavioral variables.

**Extended Data Fig. 6.**
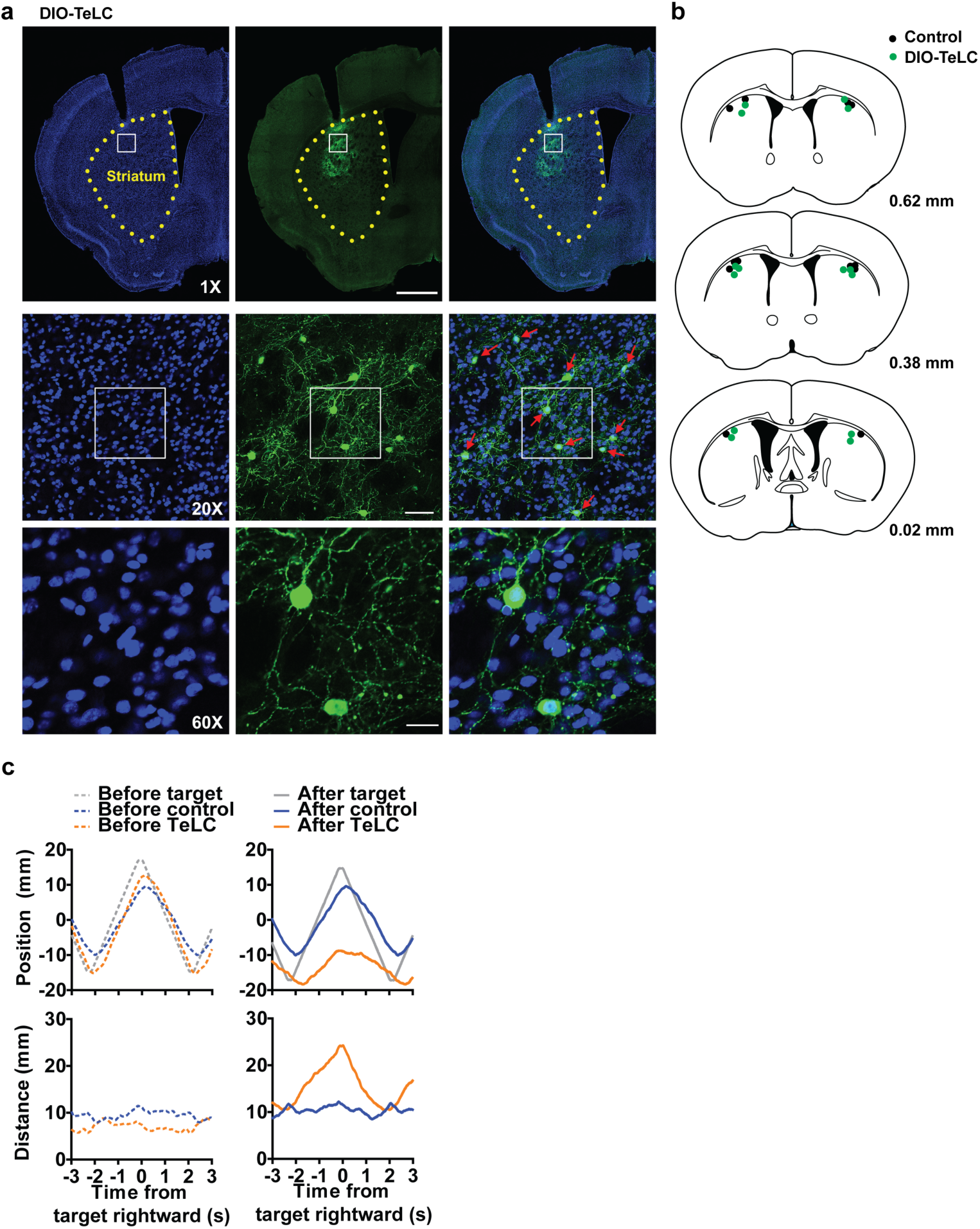
Summary of TeLC expression. a) Coronal section showing representative expression of GFP-tagged TeLC in the striatum. Left, DAPI staining. Middle, GFP staining. Right, merged. b) Locations of TeLC injection regions in the striatum. c) A representative example of target and head positions before and after TeLC injections (top). Distance error was also increased by TeLC (bottom). Both groups showed successful following behavior before TeLC injections, but silencing of FSI activity impaired pursuit performance, causing increased distance error.

**Extended Data Fig 7.**
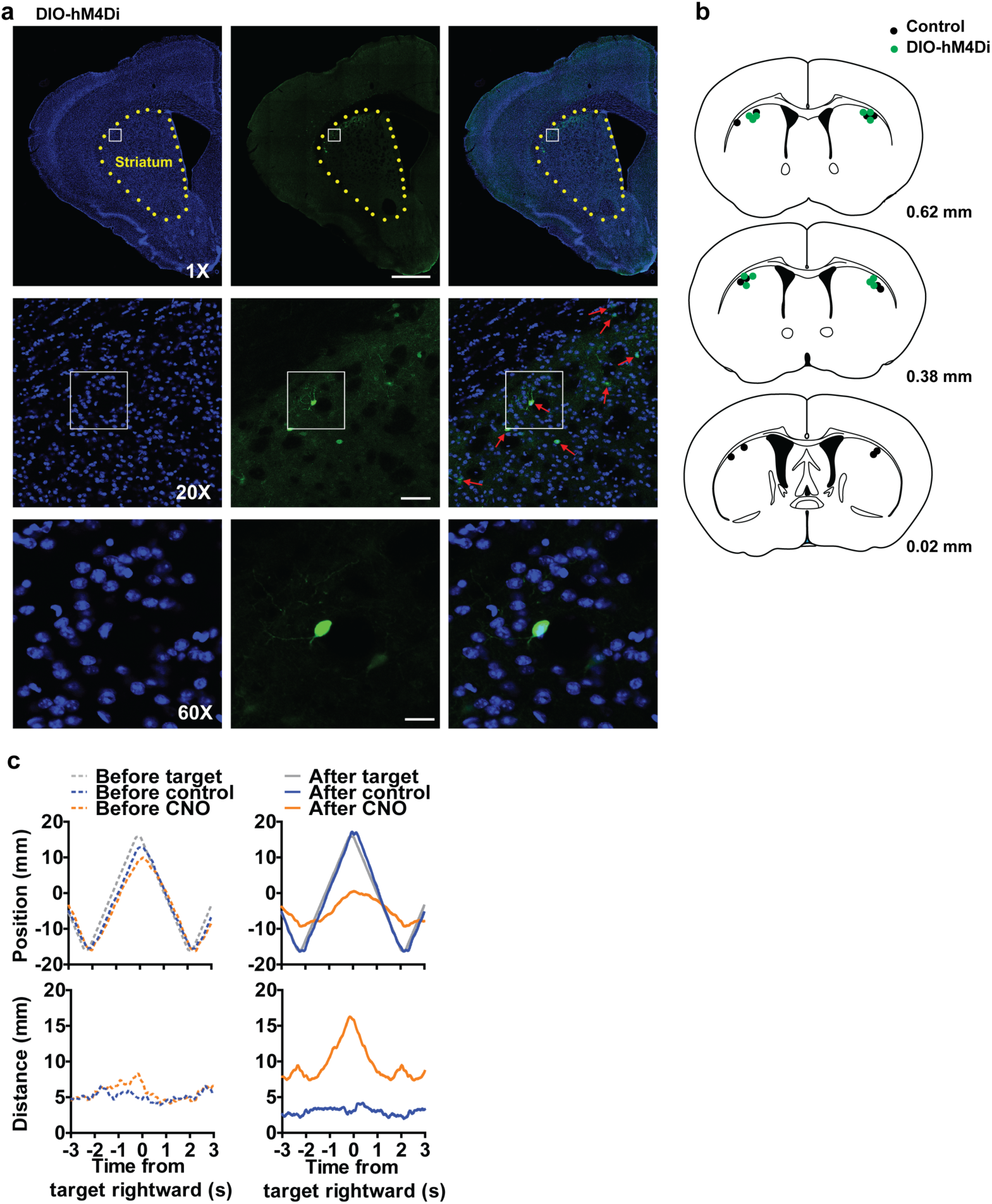
Summary of hM4Di expression. a) Coronal section showing representative expression of GFP-tagged hM4Di in the striatum. Left, DAPI staining. Middle, GFP staining. Right, merged. b) Locations of hM4Di injection regions. c) A representative example of target and head positions (control: blue; CNO hM4Di: orange) before and after CNO injection (top). Distance error was also increased by hM4Di (bottom). Both groups showed successful following behavior before CNO injections. However, blockade of FSI activity by CNO injection showed poor following behavior, causing increased distance.

**Extended Data Fig. 8.**
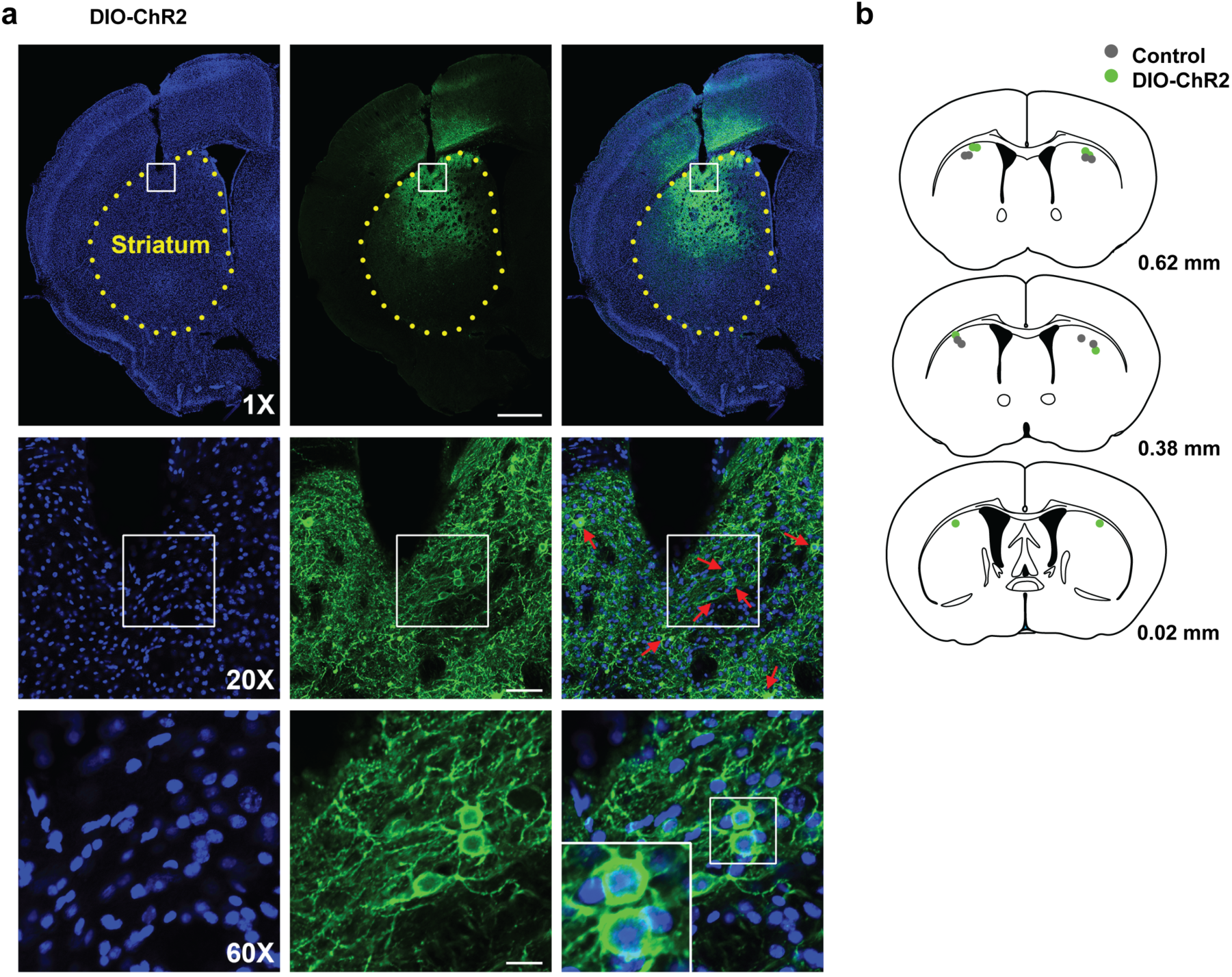
Summary of ChR2 expression. a) Coronal section showing expression of DIO-ChR2 in the sensorimotor striatum. Left, DAPI staining. Middle, GFP staining. Right, merged. *b)* Locations of optic fibers.

**Extended Data Fig. 9.**
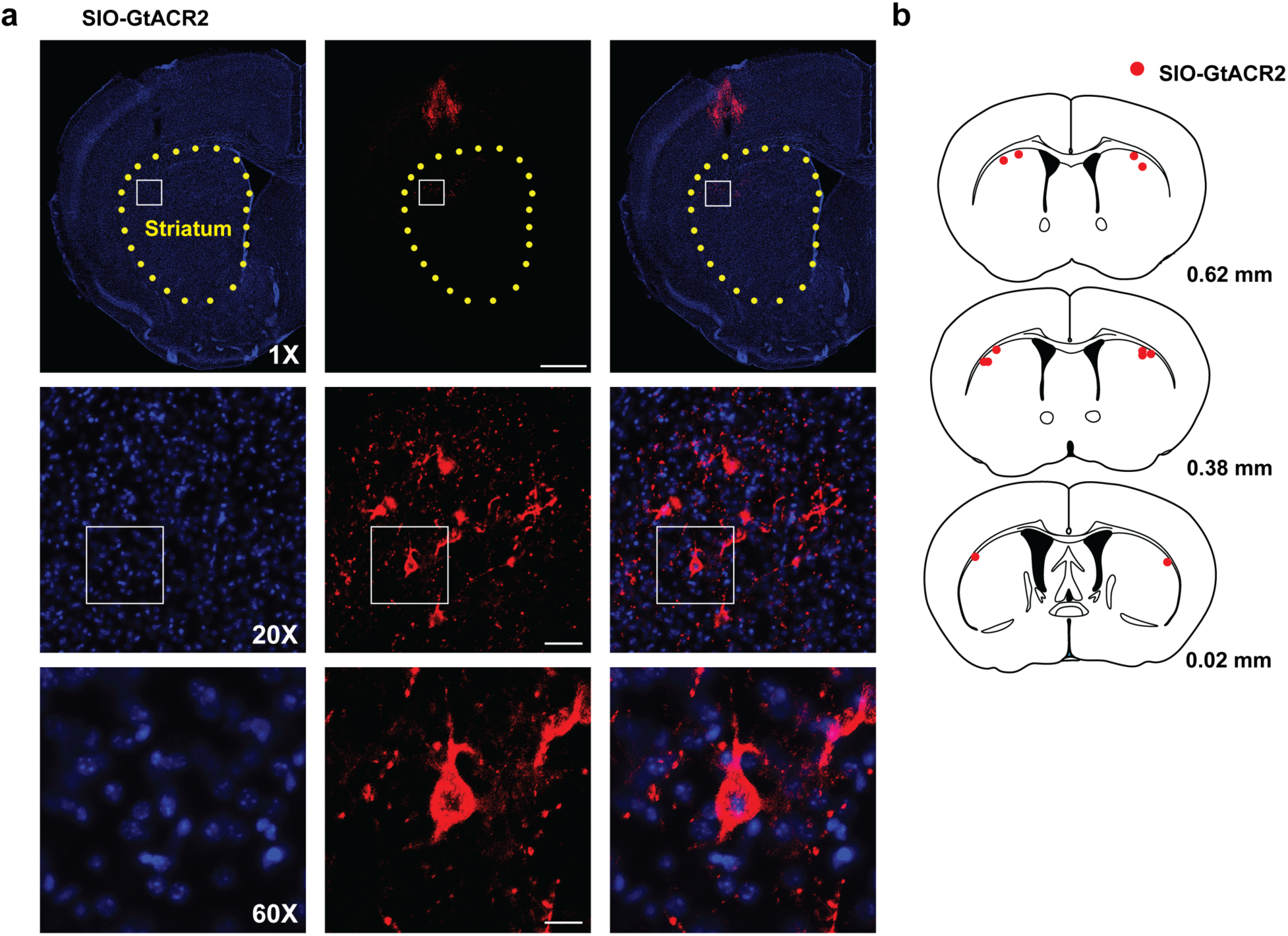
Summary of SIO-GtACR2 expression. a) Coronal section showing expression of FusionRed-tagged GtACR2 in the sensorimotor striatum. Left, DAPI staining. Middle, FusionRed staining. Right, merged. b) Locations of optic fibers.

**Extended Data Fig 10.**
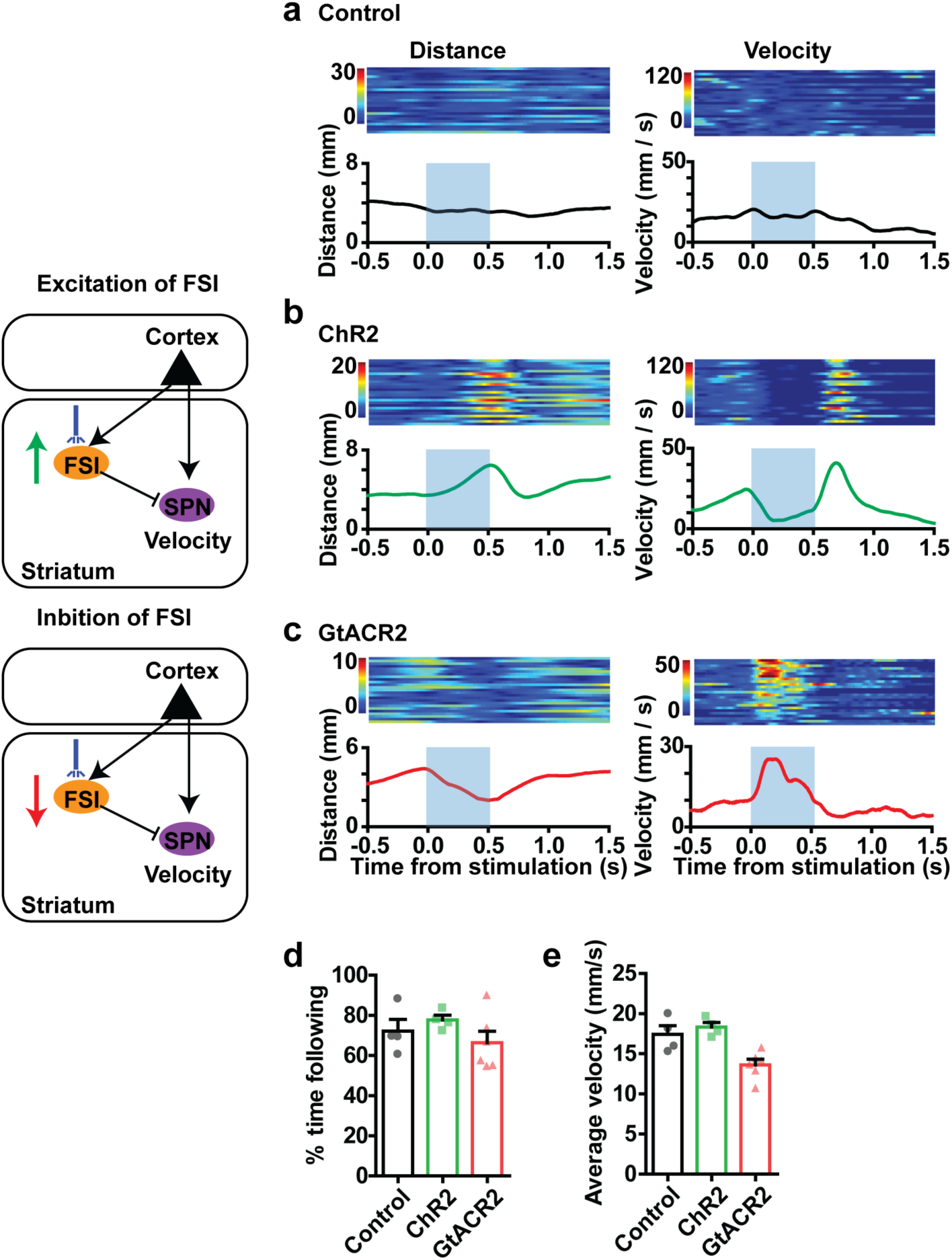
Summary of optogenetic manipulation of FSIs. *a)* Control group showed no change in distance or velocity. *b)* DIO-ChR2 group showed decreased velocity and increased distance during photo-stimulation. c) SIO-GtACR2 group showed increased velocity and decreased distance during photo-stimulation. *d)* Overall % time spent following is not altered by stimulation. e) SIO-GtACR2 group showed slightly decreased overall average velocity (p < 0.05).

**Extended Data Fig. 11.**
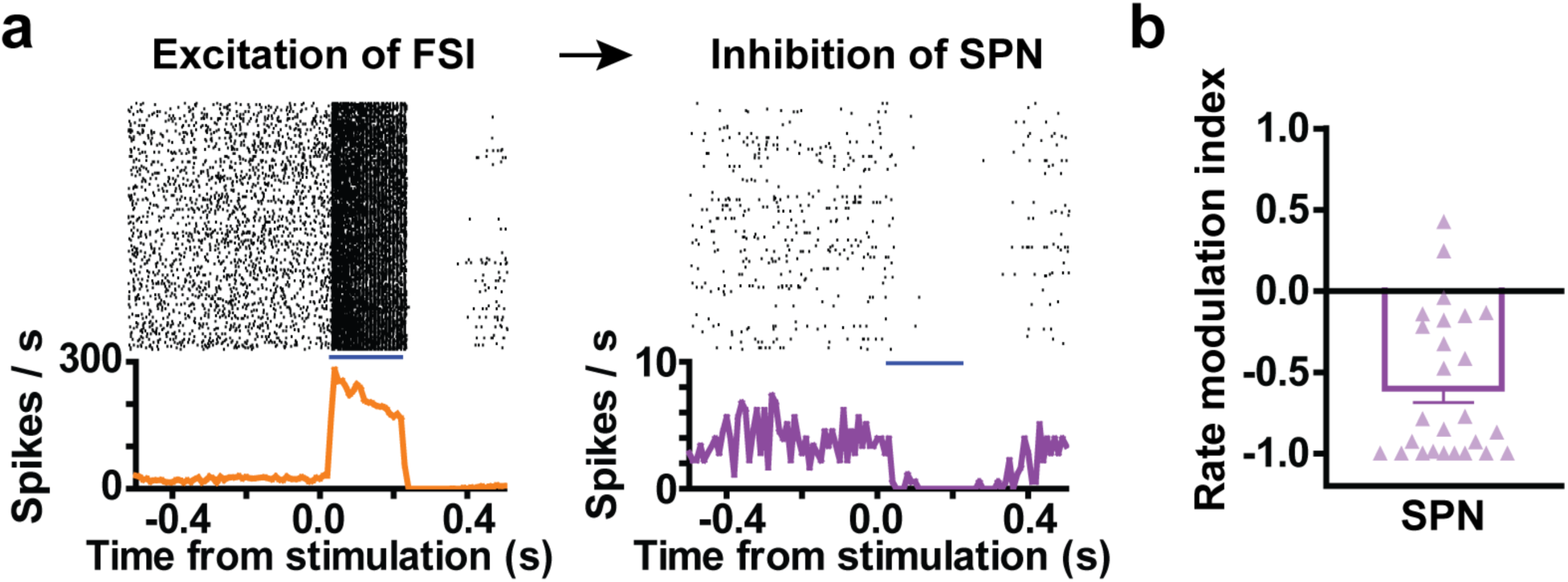
FSI-SPN interaction. a) Excitation of FSI with ChR2 *in vivo*. In the same mouse, there is disinhibition of SPNs, as shown on the right. b) Rate modulation index ((Firing rate_after_ – Firing rate_before_) / (Firing rate_after_ + Firing rate_before_)) indicates suppression of SPN activity by excitation of FSIs.

**Extended Data Fig. 12.**
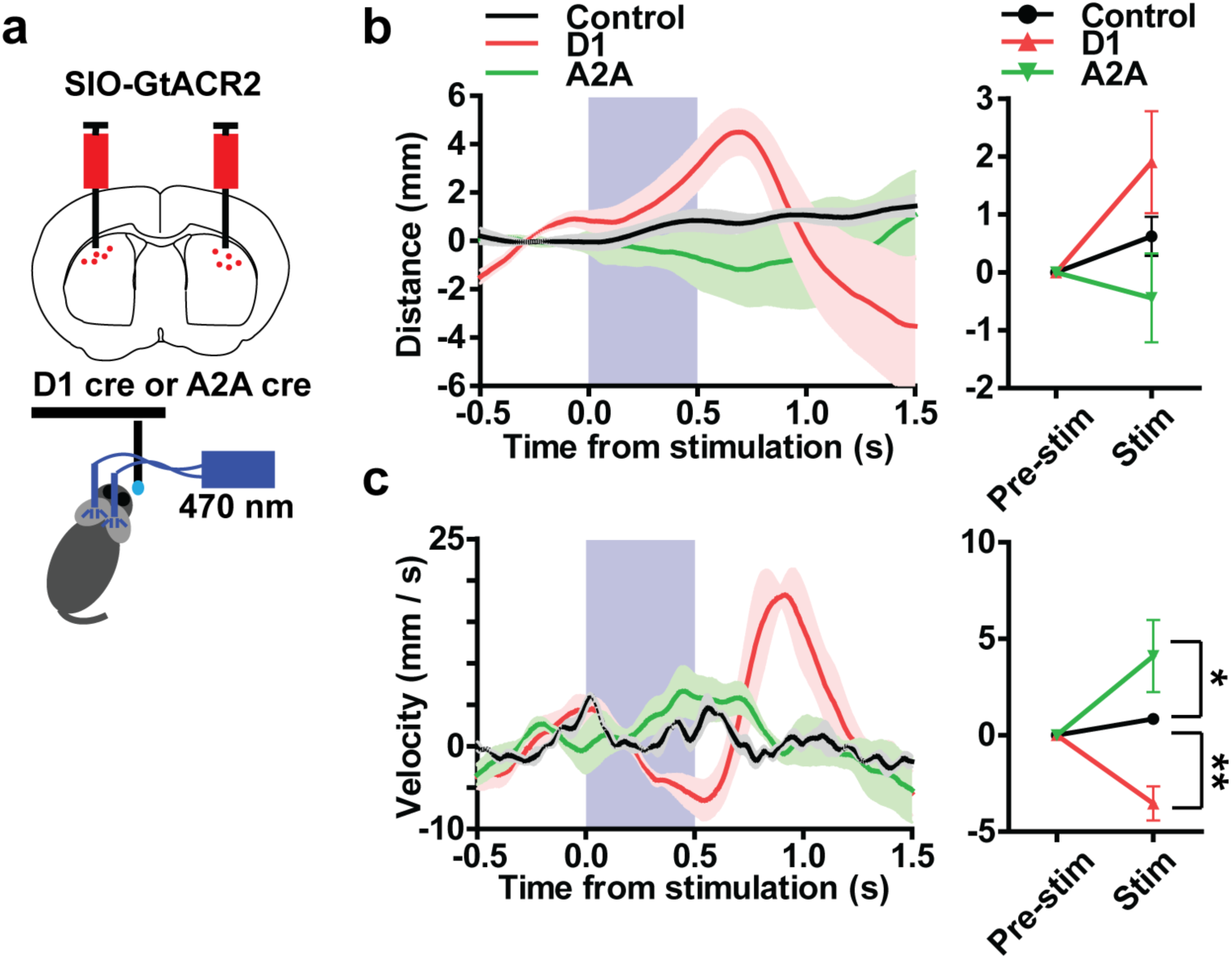
Optogenetic inhibition of SPN activity. a) We injected Cre-dependent inhibitory channelrhodopsin (AAV-hsyn1-SIO-GtACR2-Fusion Red) to reduce SPN activity in either the direct pathway (D1+ SPNs) or the indirect pathway (A2A+ SPNs). b) Effect of photo-stimulation on self-target distance. Inhibition of direct pathway or indirect pathway activity did not have a significant effect on distance (repeated measures two-way ANOVA, Interaction: F _2, 7_ = 2.484, p = 0.1530, Group: F _2, 7_ = 2.484, p = 0.1530; Time: F _1, 7_ = 2.447, p = 0.1617). Control (PV-Cre light only): n = 3; D1 SIO-GtACR2: n = 4; A2A SIO-GtACR2: n = 3. Error bar indicates ± s.e.m. c) Effect of photo-stimulation on self velocity. Inhibition of direct pathway activity decreased velocity, whereas inhibition of indirect pathway activity increased velocity (repeated measures two-way ANOVA, Interaction: F _2, 7_ = 11.82, p = 0.0057, Group: F _2, 7_ = 11.82, p = 0.0057; Time: F _1, 7_ = 0.5037, p = 0.5008, control vs D1 SIO-GtACR2, p < 0.01; control vs. A2A SIO-GtACR2, p < 0.05,). Error bar indicates ± s.e.m. ** P < 0.01. *p < 0.05.

## References

1. Krauzlis, R.J. Recasting the smooth pursuit eye movement system. J Neurophysiol 91, 591–603 (2004).

2. Lisberger, S.G., Morris, E. & Tychsen, L. Visual motion processing and sensory-motor integration for smooth pursuit eye movements. Annual review of neuroscience 10, 97–129 (1987).

3. O’Driscoll, G.A., et al. Functional neuroanatomy of smooth pursuit and predictive saccades. Neuroreport 11, 1335–1340 (2000).

4. Basso, M.A., Pokorny, J.J. & Liu, P. Activity of substantia nigra pars reticulata neurons during smooth pursuit eye movements in monkeys. European Journal of Neuroscience 22, 448–464 (2005).

5. Monteiro, P. & Feng, G. Learning from Animal Models of Obsessive-Compulsive Disorder. Biol Psychiatry (2015).

6. Yin, H.H. & Knowlton, B.J. The role of the basal ganglia in habit formation. Nature reviews. Neuroscience 7, 464–476 (2006).

7. Gerfen, C.R. & Wilson, C.J. The basal ganglia. in Handbook of chemical neuroanatomy (ed. L.W. Swanson, A. Bjorklund & T. Hokfelt) 371–468 (Elsevier, Amsterdam, 1996).

8. Mink, J.W. The basal ganglia: focused selection and inhibition of competing motor programs. Progress in neurobiology 50, 381–425 (1996).

9. Kravitz, A.V., et al. Regulation of parkinsonian motor behaviours by optogenetic control of basal ganglia circuitry. Nature 466, 622–626 (2010).

10. Gerfen, C.R., Baimbridge, K.G. & Miller, J.J. The neostriatal mosaic: compartmental distribution of calcium-binding protein and parvalbumin in the basal ganglia of the rat and monkey. Proceedings of the National Academy of Sciences 82, 8780–8784 (1985).

11. Bolam, J.P., Hanley, J.J., Booth, P.A. & Bevan, M.D. Synaptic organisation of the basal ganglia. J Anat 196, 527–542. (2000).

12. Tepper, J.M., Tecuapetla, F., Koos, T. & Ibanez-Sandoval, O. Heterogeneity and diversity of striatal GABAergic interneurons. Front Neuroanat 4, 150 (2010).

13. Gage, G.J., Stoetzner, C.R., Wiltschko, A.B. & Berke, J.D. Selective activation of striatal fast-spiking interneurons during choice execution. Neuron 67, 466–479 (2010).

14. Kalanithi, P.S., et al. Altered parvalbumin-positive neuron distribution in basal ganglia of individuals with Tourette syndrome. Proc Natl Acad Sci U S A 102, 13307–13312 (2005).

15. Burguière, E., Monteiro, P., Feng, G. & Graybiel, A.M. Optogenetic stimulation of lateral orbitofronto-striatal pathway suppresses compulsive behaviors. Science 340, 1243–1246 (2013).

16. O’Hare, J.K., et al. Striatal fast-spiking interneurons selectively modulate circuit output and are required for habitual behavior. eLife 6 (2017).

17. Kim, N., Barter, J.W., Sukharnikova, T. & Yin, H.H. Striatal firing rate reflects head movement velocity. Eur J Neurosci 40, 3481–3490 (2014).

18. Yttri, E.A. & Dudman, J.T. Opponent and bidirectional control of movement velocity in the basal ganglia. Nature 533, 402–406 (2016).

19. Bartholomew, R.A., et al. Striatonigral control of movement velocity in mice. Eur J Neurosci 43, 1097–1110 (2016).

20. Turner-Evans, D., et al. Angular velocity integration in a fly heading circuit. eLife 6 (2017).

21. Boyden, E.S., Zhang, F., Bamberg, E., Nagel, G. & Deisseroth, K. Millisecond-timescale, genetically targeted optical control of neural activity. Nature neuroscience 8, 1263–1268 (2005).

22. Ghosh, K.K., et al. Miniaturized integration of a fluorescence microscope. Nature methods 8, 871–878 (2011).

23. Cai, D.J., et al. A shared neural ensemble links distinct contextual memories encoded close in time. Nature 534, 115–118 (2016).

24. Schiavo, G., et al. Tetanus and botulinum-B neurotoxins block neurotransmitter release by proteolytic cleavage of synaptobrevin. Nature 359, 832–835 (1992).

25. Murray, A.J., et al. Parvalbumin-positive CA1 interneurons are required for spatial working but not for reference memory. Nature neuroscience 14, 297–299 (2011).

26. Armbruster, B.N., Li, X., Pausch, M.H., Herlitze, S. & Roth, B.L. Evolving the lock to fit the key to create a family of G protein-coupled receptors potently activated by an inert ligand. Proc Natl Acad Sci U S A 104, 5163–5168 (2007).

27. Zhang, F., et al. Multimodal fast optical interrogation of neural circuitry. Nature 446, 633–639 (2007).

28. Mahn, M., et al. High-efficiency optogenetic silencing with soma-targeted anion-conducting channelrhodopsins. Nature communications 9, 4125 (2018).

29. Sohal, V.S., Zhang, F., Yizhar, O. & Deisseroth, K. Parvalbumin neurons and gamma rhythms enhance cortical circuit performance. Nature 459, 698–702 (2009).

30. Zhang, F., Wang, L.P., Boyden, E.S. & Deisseroth, K. Channelrhodopsin-2 and optical control of excitable cells. Nature methods 3, 785–792 (2006).

31. Ramanathan, S., Hanley, J.J., Deniau, J.M. & Bolam, J.P. Synaptic convergence of motor and somatosensory cortical afferents onto GABAergic interneurons in the rat striatum. The Journal of neuroscience: the official journal of the Society for Neuroscience 22, 8158–8169 (2002).

32. Doig, N.M., Moss, J. & Bolam, J.P. Cortical and thalamic innervation of direct and indirect pathway medium-sized spiny neurons in mouse striatum. The Journal of neuroscience 30, 14610–14618 (2010).

33. Wilson, C.J. Basal ganglia. in The synaptic organization of the brain (ed. G.M. Shephard) (Oxford University Press, New York, 2004).

34. Barter, J.W., et al. Basal ganglia outputs map instantaneous position coordinates during behavior. Journal of Neuroscience 35, 2703–2716 (2015).

35. Yin, H.H. The Basal Ganglia in Action. Neuroscientist 23 (2017).

36. Li, Y., et al. Hypothalamic Circuits for Predation and Evasion. Neuron 97, 911–924 e915 (2018).

37. Park, S.G., et al. Medial preoptic circuit induces hunting-like actions to target objects and prey. Nature neuroscience 21, 364–372 (2018).

38. Han, W., et al. Integrated Control of Predatory Hunting by the Central Nucleus of the Amygdala. Cell 168, 311–324 e318 (2017).

39. Kirouac, G.J., Li, S. & Mabrouk, G. GABAergic projection from the ventral tegmental area and substantia nigra to the periaqueductal gray region and the dorsal raphe nucleus. The Journal of comparative neurology 469, 170–184 (2004).

40. Yin, H.H. Action, time and the basal ganglia. Philosophical Transactions of the Royal Society B: Biological Sciences 369 (2014).

41. Barter, J., et al. Beyond reward prediction errors: the role of dopamine in movement kinematics. Frontiers in Integrative Neuroscience 9, 39 (2015).

42. Barter, J.W., Castro, S., Sukharnikova, T., Rossi, M.A. & Yin, H.H. The role of the substantia nigra in posture control. European Journal of Neuroscience 39 (9), 1465–1473 (2014).

43. Fan, D., Rossi, M.A. & Yin, H.H. Mechanisms of action selection and timing in substantia nigra neurons. The Journal of neuroscience: the official journal of the Society for Neuroscience 32, 5534–5548 (2012).

44. Yin, H.H. How basal ganglia outputs generate behavior. Advances in neuroscience 2014, 768313 (2014).

45. Yin, H.H. The role of opponent basal ganglia outputs in behavior. Future Neurology 11, 149–169 (2016).

46. Bracci, E. & Panzeri, S. Excitatory GABAergic effects in striatal projection neurons. J Neurophysiol 95, 1285–1290 (2006).

47. Isomura, Y., et al. Reward-modulated motor information in identified striatum neurons. The Journal of Neuroscience 33, 10209–10220 (2013).

48. Owen, S.F., Berke, J.D. & Kreitzer, A.C. Fast-Spiking Interneurons Supply Feedforward Control of Bursting, Calcium, and Plasticity for Efficient Learning. Cell 172, 683–695 e615 (2018).

49. Rueda-Orozco, P.E. & Robbe, D. The striatum multiplexes contextual and kinematic information to constrain motor habits execution. Nature neuroscience (2015).

50. Klaus, A., et al. The Spatiotemporal Organization of the Striatum Encodes Action Space. Neuron 95, 1171–1180 e1177 (2017).

51. Turner, R.S. & Desmurget, M. Basal ganglia contributions to motor control: a vigorous tutor. Current opinion in neurobiology 20, 704–716 (2010).

52. Georgopoulos, A.P., DeLong, M.R. & Crutcher, M.D. Relations between parameters of step-tracking movements and single cell discharge in the globus pallidus and subthalamic nucleus of the behaving monkey. The Journal of Neuroscience 3, 1586–1598 (1983).

53. Gritton, H.J., et al. Unique contributions of parvalbumin and cholinergic interneurons in organizing striatal networks during movement. Nature neuroscience (2019).

54. Cui, G., et al. Concurrent activation of striatal direct and indirect pathways during action initiation. Nature 494, 238–242 (2013).

55. Rieke, F. Spikes: exploring the neural code (The MIT Press, 1999).

56. Geddes, C.E., Li, H. & Jin, X. Optogenetic Editing Reveals the Hierarchical Organization of Learned Action Sequences. Cell 174, 32–43 e15 (2018).

57. Ade, K.K., Wan, Y., Chen, M., Gloss, B. & Calakos, N. An Improved BAC Transgenic Fluorescent Reporter Line for Sensitive and Specific Identification of Striatonigral Medium Spiny Neurons. Front Syst Neurosci 5, 32 (2011).

58. Rossi, M.A., Fan, D., Barter, J.W. & Yin, H.H. Bidirectional Modulation of Substantia Nigra Activity by Motivational State. PloS one 8, e71598 (2013).

59. Fan, D., et al. A wireless multi-channel recording system for freely behaving mice and rats. PLoS One 6, e22033 (2011).

60. Zhou, P., et al. Efficient and accurate extraction of in vivo calcium signals from microendoscopic video data. eLife 7 (2018).

61. Pnevmatikakis, E.A., et al. Simultaneous Denoising, Deconvolution, and Demixing of Calcium Imaging Data. Neuron 89, 285–299 (2016).

62. Paxinos, G. & Franklin, K. The mouse brain in stereotaxic coordinates (Academic Press, New York, 2003).

63. Sparta, D.R., et al. Construction of implantable optical fibers for long-term optogenetic manipulation of neural circuits. Nature protocols 7, 12–23 (2012).

